# The axes of biology: a novel axes-based network embedding paradigm to decipher the functional mechanisms of the cell

**DOI:** 10.1101/2023.07.31.551263

**Authors:** Sergio Doria-Belenguer, Alexandros Xenos, Gaia Ceddia, Noël Malod-Dognin, Nataša Pržulj

## Abstract

Common approaches for deciphering biological networks involve network embedding algorithms. These approaches strictly focus on clustering the genes’ embedding vectors and interpreting such clusters to reveal the hidden information of the networks. However, the difficulty in interpreting the genes’ clusters and the limitations of the functional annotations’ resources hinder the identification of the currently unknown cell’s functioning mechanisms. Thus, we propose a new approach that shifts this functional exploration from the embedding vectors of genes in space to the axes of the space itself. Our methodology better disentangles biological information from the embedding space than the classic gene-centric approach. Moreover, it uncovers new data-driven functional interactions that are unregistered in the functional ontologies, but biologically coherent. Furthermore, we exploit these interactions to define new higher-level annotations that we term Axes-Specific Functional Annotations and validate them through literature curation. Finally, we leverage our methodology to discover evolutionary connections between cellular functions and the evolution of species.

## Introduction

Cells are the basic building blocks of all living organisms. Understanding the complex intracellular processes is crucial not only for identifying the fundamental mechanisms of life, but also for deciphering the evolutionary history of species. Advances in capturing technologies have yielded a massive production of large-scale molecular data that describe the complex machinery of the cell [1]. These data are often modeled as networks, in which nodes are molecular entities and the edges connecting them represent their relationships: e.g., in protein-protein interaction networks (PPIs), nodes represent proteins and edges indicate physical interactions (bindings) between them, as measured by biological experiments. These networks are a valuable source of biological information that need to be “untangled” by new algorithms to reveal the information hidden in their wiring patterns.

Recent approaches for deciphering these complex data are based on network embedding techniques [2, 3]. These algorithms aim to find the vectorial representations of the network nodes in a low-dimensional embedding space spanned by a system of coordinates (a.k.a., embedding axes) while preserving the structural information of the network [2, 4]. Defining an optimal number of dimensions of the embedding space is key to properly capturing the structural information of the network. However, there is no gold-standard approach to finding the optimal dimensionality of the embedding space, so researchers rely on grid search, domain knowledge or heuristics [5].

Network embedding techniques include different algorithms, such as Natural Language Processing (NLP)-inspired methods based on Neural Networks (NNs), e.g., DeepWalk [6], LINE [7] and node2vec [8]. Other approaches include matrix factorization-based ones, including e.g.: Singular Value Decomposition (SVD) [9], Principal Component Analysis (PCA) [10] with the sparse and probabilistic variants [11], Independent Component Analysis (ICA) [12] and Non-negative Matrix Factorizations (NMF) [13, 14]. In particular, Non-negative Matrix Tri-Factorization (NMTF) is an extension of NMF and a well-known Machine Learning (ML) technique introduced for co-clustering and dimensionality reduction [15]. Contrary to the NLP-inspired methods, the embedding spaces produced by NMTF have valuable properties, e.g., orthonormality and non-negativity. It has been hypothesized that such properties may lead to an easier interpretation and deeper scientific insight [16]. However, this hypothesis has not been tested, so the real impact of these properties remains unclear. Revealing the information in the complexity of a molecular network requires not only an embedding algorithm, but also methods to translate the results into biologically interpretable models [17]. Gene-centric embedding approaches analyze the topological and functional properties of molecular networks by clustering together genes whose embedding vectors are in proximity in the embedding space. These clusters represent subgraphs of the molecular network that display significant clustering properties, i.e., genes within each cluster are more densely connected to each other than to genes outside the cluster. To functionally interpret these clusters, current methods rely on several curated ontologies, such as the Kyoto Encyclopedia of Genes and Genomes (KEGG) [18], Reactome [19] and the Gene Ontology (GO) [20]. Among them, GO has the largest number of records [20, 21]. GO terms are often used in functional enrichment analysis to evaluate the statistical over-representation of biological functions in genes’ clusters [22]. These statistically enriched functions are then used to characterize the cell’s functional organization from a molecular network captured by the gene clusters produced by embeddings of genes [23].

Gene-centric methods have demonstrated their potential in functionally explaining molecular networks, leading to a better understanding of the cells’ machinery [23, 24]. However, they present several limitations that hinder the identification of the functional mechanisms of the cells. First, they rely on statistically overrepresented genes’ functions (described by GO terms), which only represent a subset of all the available GO terms. Second, the genes’ clusters often present redundant enriched GO terms [24], further reducing the functional information uncovered.

Recently, we introduced the Functional Mapping Matrix (FMM) as a new approach to overcome the limitations of the gene-centric embedding approaches [25]. The FMM uncovers the cell’s functional organization by capturing the functional interactions between all GO Biological Process (BP) terms based on their mutual positions in the embedding space. Unlike gene-centric approaches, the FMM uses all available GO BP terms to generate a complete functional map of the cell’s organization. However, while our FMM captures all pairwise interactions between GO terms, it does not allow for identifying the most relevant ones for biological interpretation. Additionally, similar to gene-centric methods, the FMM is limited by current ontologies: the analysis is limited to the predefined set of functional annotations, i.e., new functions that are not described in the ontology can not be uncovered. Additionally, due to the incompleteness of the ontologies [17, 21], no functional information can be captured from unannotated parts of the network. Finally, the slow update rate of annotations by database curators poses a bottleneck, limiting the practical use of these annotations [17, 21].

To overcome these limitations, we propose to use the axes of the embedding space to identify the functional mechanisms of a cell. Contrary to the current state-of-the-art approaches that focus on the organization of the embedded entities (genes and their functions) in the embedding space, our method focuses on the axes of the embedding space itself to define higher-order processes that are not described in the current ontologies and represent important cellular mechanisms. To this end, we generate the gene embedding spaces of six species, *Homo sapiens sapiens* (human), *Saccharomyces cerevisiae* (budding yeast), *Schizosaccharomyces pombe*(fission yeast), *Rattus norvegicus* (rat), *Drosophila melanogaster* (fruit fly) and *Mus musculus* (mouse), by applying the NMTF and Deepwalk NN-based algorithm to the corresponding species PPI networks. We apply the NMTF algorithm with and without orthonormality constraints (“ONMTF” and “NMTF,” respectively) to gain insights into their impact on the functional organization of the embedding space axes. Then, to untangle the biological information hidden in the resulting gene embedding spaces, we embed GO BP terms in these spaces and associate them with the axes of each space as described below.

We demonstrate that the axes of the embedding space better disentangle biological information from the embedding space than gene-centric clustering followed by functional enrichment analysis: with semantically similar GO BP terms get associated with the same axis, i.e., each axis represents a specific biological function. Moreover, we show that the axes of the ONMTF gene embedding spaces better untangle biological information from the embedding space than Deepwalk and that this information is more coherently stratified across the axes. We demonstrate that this observation is connected to the properties of the ONMTF embedding spaces, such as orthonormality and non-negativity, which improve the organization of such embedding spaces.

Furthermore, we use our novel axes-based method to uncover the optimal dimensionality of the different species PPI network embedding spaces. For the optimal dimensionality, we explore the meaning of the GO BP terms associated with their axes. We observe that while the GO BP terms that are associated with the same axis tend to be functionally related (with large semantic similarity), our axes also associate seemingly unrelated GO-BP terms. We validate these new data-driven interactions by literature curation and show that they are biologically coherent and represent the functional interactions between GO BP terms in higher-order cellular functions, e.g., cell adhesion processes. Moreover, our methodology predicts new functional interactions for which we find some literature validation, but whose interaction has yet to be experimentally validated, e.g., the role of ribosomal RNA transcription in midbrain development.

Also, we investigate the higher-order cellular functions that raise from the GO BP terms’ functional interactions captured by the axes. To this end, we summarize all the GO terms that are associated with a given axis into a higher-level functional annotation that we term ASFA (Axes-Specific Functional Annotation). We find that ASFAs not only define coherent biological processes, such as the cellular response to misfolded proteins, but also they can be exploited to find new evolutionary connections between species, e.g., the mechanisms behind neural synapses that were inherited from prokaryotic organisms.

Finally, due to the incompleteness of GO annotations, we find that not all axes have associated GO terms, i.e., the biological meaning of the non-annotated axes can not be discovered using the current functional annotations. We go beyond this limitation and propose to use the description of the genes that are associated with the axes to define the ASFAs. We demonstrate that the corresponding ASFAs are biologically coherent and complement ASFAs obtained from the GO BP terms associated with the axes.

## Results

### The axes of the embedding spaces capture the cell’s functional organization

In this section, we evaluate if the axes of the embedding space of PPI networks uncover the cell’s functional organization. To this end, we generate the embedding spaces of six species (human, budding yeast, fission yeast, rat, fruit fly, and mouse) by applying ONMTF, NMTF, and Deepwalk algorithms on the corresponding species PPI networks (detailed in Methods, sections Biological datasets and Embedding the PPI networks). To analyze the impact of the embedding space’s dimensionality on the ability of the embedding methods to reveal the cell’s functional organization, for each species PPI network and for each embedding method, we generate the embedding spaces with increasing dimensionalities (from 50 to 1,000 dimensions with a step of 50). Then, we embed GO BP terms into these embedding spaces and use the axes of the embedding space to capture the embedded GO BP terms (detailed in Methods, section Annotating the axes of the gene embedding space with GO BP terms). We evaluate the ability of the embedding space axes to uncover the cell’s functional organization by analyzing the percentage of axes having at least one associated GO BP term, the percentage of the total GO BP terms that are associated with the axes, and the functional similarity of the captured GO BP terms (detailed in Methods, section Assessing the ability of our axes-based methodology to capture biological information). Here, we detail the results of the human PPI network and comment on whether similar results hold for the other species.

We observe that Deepwalk embedding spaces have, on average, the largest number of axes with associated GO BP terms (88.05%) followed by ONMTF (59.25%) and NMTF (42.78%), see Figure 1. However, the axes of ONMTF embedding spaces capture a larger number of GO BP terms (37.12%) than the axes of Deepwalk (33.8%), and NMTF embedding spaces (11.95%), see Figure 1. These results suggest that Deepwalk embedding spaces capture fewer biological functions, but “spread” them more across the axes (average of 9.7 GO BP terms per axis), while ONMTF spaces capture more biological functions and group them on a smaller number of axes (average of 16 GO BP terms per axis). Furthermore, we investigate whether this captured information is coherently stratified across the axes, i.e., if the GO BP terms that are associated with the same axis are more functionally similar (higher semantic similarity and closer in the ontology’s directed acyclic graph, DAG) than those associated with different axes (detailed in Methods, section Assessing the ability of our axes-based methodology to capture biological information). We find that ONMTF embedding spaces, not only group more GO BP terms per axis, but the functions that are associated with the same axis are functionally more coherent (3.12 times higher average semantic similarity than expected by random, Mann-Whitney U test with p-value 3.39 × 10^−8^) than the ones associated with the axis in NMTF and Deepwalk embedding spaces (2.6 and 2.1 times larger than expected by random, respectively, Mann-Whitney U test with p-values 3.38 × 10^−8^ and 2.41 × 10^−7^, respectively), see Supplementary Table 4. Moreover, GO BP terms associated with the axes of the ONMTF spaces are on average closer in the ontology DAG (average shortest path of 4.21), than the ones captured in NMTF and Deepwalk embedding spaces (average shortest path of 4.70, and 5.35, respectively). We find similar results for the rest of the species embedding spaces (see Supplementary Tables 3 and 5).

**Fig. 1:**
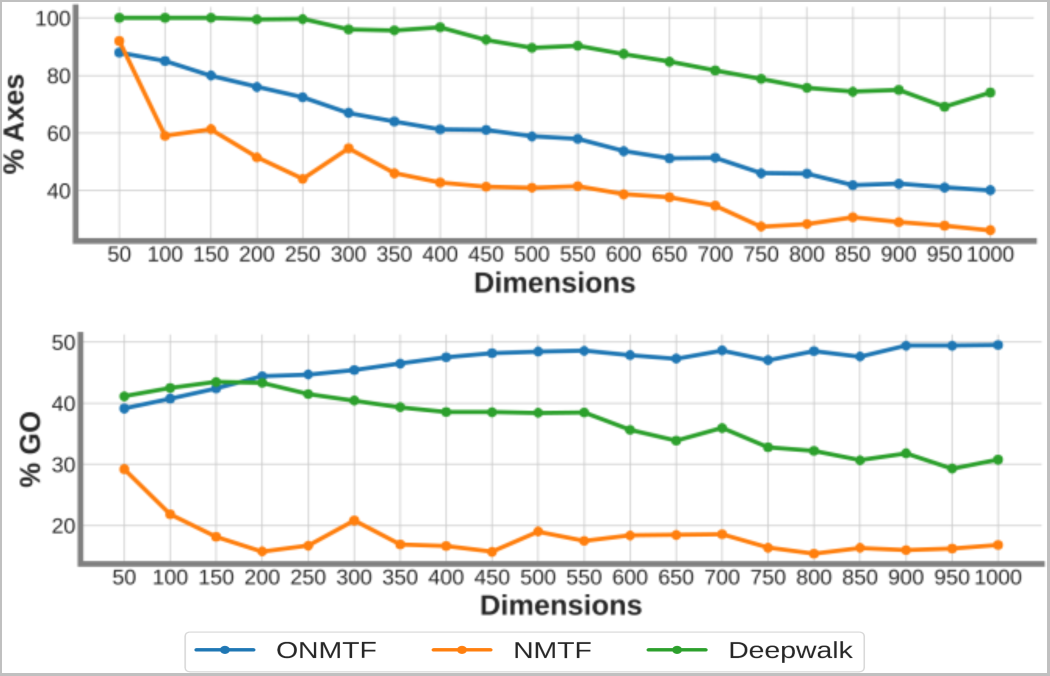
The axes of the human ONMTF, NMTF, and Deepwalk embedding spaces uncover the cell’s functional organization from the human PPI network. The top panel shows the percentage of axes that capture at least one embedded GO BP term. The bottom panel shows the percentage of the total embedded GO BP terms that are captured by the axes of the space. For each panel, the horizontal axis displays the number of dimensions of the embedding space. For each panel, the color of the lines corresponds to the three tested embedding algorithms: ONMTF (blue), NMTF (orange) and Deepwalk (green).

These results confirm that the embedding axes capture biological knowledge from the PPI network and that this information is biologically meaningfully distributed across dimensions, i.e., each axis captures a set of GO terms that are functionally related. Moreover, we demonstrate that the axes of the ONMTF embedding spaces capture more and better-stratified functional information than the other methods. We show that this ability of ONMTF to produce embedding spaces whose axes capture more and better stratified functional information can be attributed to the properties of the embedding spaces produced by ONMTF, i.e., orthonormality and non-negativity (Supplementary Results, section Orthonormality and positive constraints improve the functional organization of the gene embedding space). Thus, embedding in positive and orthonormal spaces, which only NMTF-based frameworks allow for, leads to the embedding spaces that best capture the cell’s functional organization from the biological networks. In addition, we find that the embedding space’s dimensionality also affects the biological information captured by the embedding space axes, e.g. the total amount of GO BP terms captured flattens after 400-500 dimensions in ONMTF embedding spaces (see Figure 1). This indicates that increasing the embedding space’s dimensionality enables better disentanglement of functions encoded in the PPI networks, but that adding more dimensions beyond 500 does not improve the capture of either more biological information, or more specific information from the ONMTF embedding space (Supplementary Results, section Exploring the impact of the network embedding space’s dimensionality on the biological information captured by the axes). Thus, we choose 500 dimensions as the optimal dimensionality for the human ONMTF embedding space (the optimal dimensionality for other species embedding spaces is included in Supplementary Table 6). This optimal dimensionality is coherent with the number of dimensions usually applied in NLP [26, 27].

Also, we compare the ability of our axes-based method to uncover functional information from PPI network embedding spaces with the standard gene-centric approach. To this end, we cluster genes based on the cosine similarity of their embedding vector and apply functional enrichment analysis to identify the GO BP terms that are enriched in the resulting genes’ clusters (Supplementary Results, section Our axes-based method outperforms the classic gene-centric approach in capturing the cell’s functional organization). In particular, our axes-based methodology captures 1.32 times more functional information (GO BP terms associated with the axes) from the ONMTF embedding spaces than the standard gene-centric approach (GO BP terms enriched in at least one gene cluster). Moreover, our axes-based methodology better stratifies the information captured, as GO BP terms associated with the same axis are, on average, 1.42 times more semantically similar than those enriched in the same gene cluster.

Furthermore, we compare our axes-based methodology to our function-centric approach based on FMM [25], which measures the association between functional annotations based on the cosine distances of their embedding vectors in the embedding space of a network (Supplementary Results, section Our axes-based method is in agreement with the FMM-based methodology). We find that the functional information captured by our axes-based methodology is significantly in agreement with our previous FMM, i.e., pairs of GO BP terms associated with the same axis are located close in the embedding space having small association values in the FMM (Mann-Whitney U test with p-value ≤ 1 × 10^−323^ for ONMTF embedding spaces). However, unlike FMM, our new methodology enables the identification of the most relevant functional interactions between GO BP terms for biological interpretation. These results suggest that exploiting the axes of the embedding space itself better disentangles functional information from the PPI embedding spaces compared to relying solely on the organization of embedded entities (genes and gene functions) within the space.

In conclusion, we show that the axes of the ONMTF embedding spaces better uncover the cell’s functional organization and define the optimal dimensionality of such spaces. In the following sections, we investigate the biological meaning of the axes of the optimally dimensional ONMTF embedding spaces of PPI networks of different species.

### The axes of the embedding space uncover new functional interactions between GO BP terms

We investigate whether our axes-based methodology captures new data-driven interactions between GO BP terms that are not described in the Gene Ontology, but are biologically coherent. Recall that interactions between GO terms in the ontology reflect their functional similarity (semantic similarity) [28]. Thus, first we explore if seemingly unrelated GO BP terms are associated with the same axis. To this end, we compute Lin’s semantic similarity between all two GO BP terms (detailed in Methods, section Quantifying the evolutionary conservation of biological functions). Then, for each axis, we take the average semantic similarity among its pairs of associated GO BP terms (“intra-axis SeSi”). By taking all the “intra-axis SeSi” over all the embedding axes, we define the distribution of “intra-axis SeSi” (see the distribution in Supplementary Figure 6).

Over all species PPI network embedding spaces, we find an average “intra-axis SeSi” of 0.52, ranging from 0.99 down to 0.10. Thus, while GO BP terms associated with the same axis tend to be functionally related, our axes also associate seemingly unrelated GO BP terms (according to GO). We further assess the biological coherence of associating unrelated GO BP terms with the same axis by conducting a literature curation. For this analysis, we focus on the embedding axes that capture the highest number of seemingly unrelated GO BP terms, i.e., those with a significantly low “intra-axis SeSi” (detailed in the Supplementary Results, section The axes of the embedding space uncover new functional interactions between GO BP terms in different model organisms). For each of these axes, we evaluate if the interactions between its associated GO BP terms are biologically coherent.

Among the 13 axes with a significantly low “intra-axis SeSi” in the human PPI network embedding space, we find 7 axes (53.8%) for which all captured functional interactions are known in the literature to occur in humans, 3 axes (23.1%) for which captured functional interactions are described in the literature to occur in model organisms, but not yet in human and 3 axes (23.1%) for which captured functional interactions are not described in the literature, but are yet biologically coherent from a functional perspective.

One example of an axis that captures known functional interactions in human is axis 37, which has two associated GO BP terms, namely GO:0033624 (negative regulation of integrin activation) and GO:0032487 (regulation of Rap protein signal transduction). Although these functions are not connected based on the Gene Ontology (semantic similarity of 0.16), their functional interaction is biologically coherent, since Rap proteins are known to be involved in integrin-mediated cell adhesion in humans [29, 30].

On the other hand, axis 59 is an example of an axis that captures functional interactions only described in model organisms. It has two associated GO BP terms: GO:0044528 (regulation of mitochondrial mRNA stability) and GO:0006891 (intra-Golgi vesicle-mediated transport). These functions are not connected based on the Gene Ontology (semantic similarity of 0.08). However, Gerards *et. al* [31] demonstrated that vesicle transport at the Golgi apparatus is fundamental for the mitochondrial quality control of fruit flies. This quality control includes several processes, such as the mitochondrial unfolded protein response, which are highly influenced by mitochondrial mRNA stability[32].

Finally, axis 116 (represented in Figure 2) exemplifies an axis capturing new functional interactions. It has four associated GO BP terms. Three of them are GO:0006361 (transcription initiation at RNA polymerase I promoter), GO:0036369 (transcription factor catabolic process), and GO:1905524 (negative regulation of protein autoubiquitination). Although these terms are not connected based on the Gene Ontology (average semantic similarity of 0.29), their interaction is functionally coherent. The attachment of ubiquitin initiates the transcription factor catabolic process [33], which limits the availability of these proteins and their binding to genomic promoters like RNA polymerase I promoters [34]. Genes with RNA polymerase I promoters encode ribosomal RNAs (rRNA) crucial for ribosome construction. Hence, these three GO BP terms participate in the regulation of rRNA transcription. In contrast, the fourth GO BP term associated with axis 116 (GO:0030901) is highly semantically dissimilar to the previous three (semantic similarity of 0.05) and represents midbrain development. The dysregulation of rRNA transcription has been implicated in various neurological pathologies, including Parkinson’s disease [35] and Alzheimer’s disease [36], underscoring the significance of this regulatory process in brain function. However, to date, the involvement of rRNA synthesis in the early developmental stages of the brain remains unexplored.

**Fig. 2:**
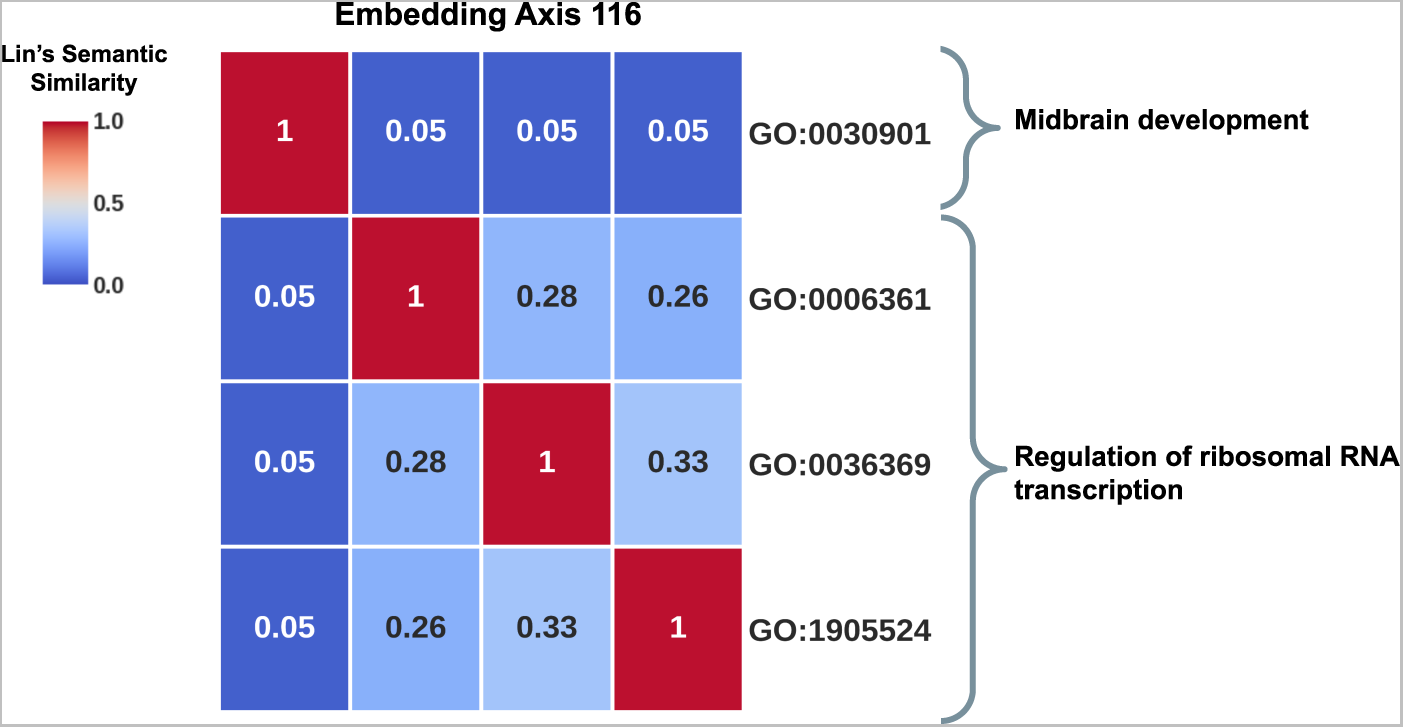
The embedding axes uncover new functional interactions between cellular functions. The plot shows the pairwise Lin’s semantic similarity between the GO BP terms associated with the axis 116 of the human PPI network embedding space. As shown in the plot, three GO BP terms associated with this axis participate in the regulation of ribosomal RNA transcription. In contrast, GO:0030901, which represents midbrain development, is not related to this group of GO BP terms according to the Gene Ontology.

In conclusion, the functional interactions captured by the axes are in good agreement with GO ontology. However, we show that the axes also captured new, data-driven interactions not described in GO that represent higher-order cellular processes (see Supplementary Results, section The axes of the embedding space uncover new functional interactions between GO BP terms in different model organisms, for an extended discussion). In the next section, we investigate the higher-order cellular processes that arise from the functional interactions captured by our methodology.

### The axes of the embedding space capture the functional mechanisms of the cell

We analyze the higher-order cellular processes that arise from the functional interactions between the GO BP terms captured by each embedding axis. To this end, we summarize the set of GO BP terms captured by each axis into ASFAs. ASFAs are composed of a set of keywords that provide a synopsis of the annotations associated with the axes (detailed in Methods, section Generating the Axes-Specific Functional Annotations). We assess whether ASFAs correctly summarize the set of GO BP terms captured by the axes and evaluate their coherence in describing biological functions through literature curation.

We find that ASFAs correctly summarize the biological information captured by the axes and describe coherent human cellular functions (see Table 1). For instance, axis 51 captures twenty-seven GO BP terms that describe the regulation of telomere (GO:0032204, GO:0032206, GO:0032210, GO:0032212, GO:1904356, GO:1904358, GO:0051972, GO:0051973, GO:1904816, GO:1904851, GO:2001252, 0070203 and GO:1904814), chromosome stability (GO:0033044), Cajal body (GO:0090666, GO:0090670, GO:1904872 and GO:1904874), protein location and stability (GO:0031647, GO:0050821, GO:0070202, GO:1903829 and GO:1904951), and RNA location to the nucleus and gene expression (GO:0006403, GO:2000573, GO:0090685 and GO:2000278). The resulting ASFA combines and summarizes the keywords of these terms (maintenance, activity, telomere, scaRNA, RNA, telomeric, biosynthetic, Cajal, regulation, protein, stability, nucleus, localization, process, lengthening, establishment, body, DNA, positive, via, stabilization, telomerase, chromosome and organization) and reflects the functions of Cajal bodies, including biogenesis and modification of Cajal body-specific RNPs (scaRNPs) and telomerase [37].

**Table 1:**
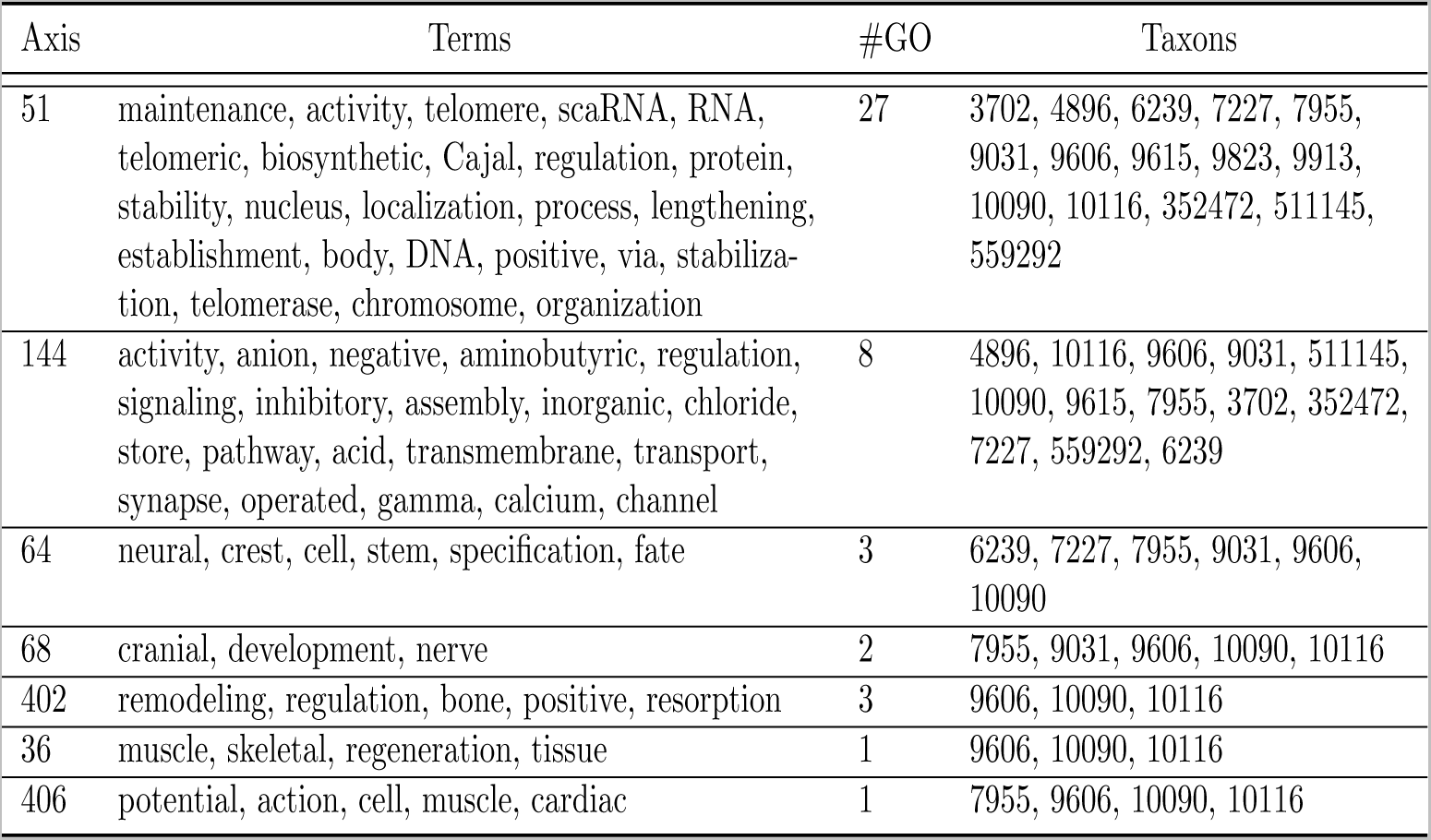
The ASFAs describe coherent functions of the human cell. For the human ONMTF embedding space, we use the GO BP terms associated with its axes to generate the ASFAs. The first column, “Axis,” lists the number of the axes from which each ASFA was obtained. The second column, “Terms,” shows the description of the ASFAs. The third column, “#GO,” displays the number of GO BP terms that are associated with the axis. The fourth column, “Taxons,” shows the Taxonomy ID of the different species for which the associated GO BP terms appear. The complete Table with all the human ASFAs and assigned GO terms can be found in the Supplementary online data.

The observed biological coherence of our ASFAs can be explained by the fact that GO BP terms associated with the axes are already functionally coherent (see Supplementary Figure 6). We find similar results for the other species (see Supplementary Results, section The Axes of the embedding space synthesize the core functions of different species’ cells, for more examples of human and the other species).

So far, we have used GO BP terms to find the higher-order functions represented by the axes of the embedding space. However, due to the incompleteness of GO annotations (only 48.6% of human genes in the PPI network are annotated), not all axes have GO BP terms associated with them (41.2% of the ONMTF human embedding space’s axes remain unannotated). Thus, we ask if these unannotated axes still carry functional information (Supplementary Results, section Non-Annotated Axes also capture the functional mechanisms of the cell). To this end, we propose to associate genes with the embedding space’s axes and use the text descriptions of the genes from the Alliance of Genome Resources database to generate the corresponding ASFAs. Using this approach, we obtain the ASFAs for 97.8% of the ONMTF human embedding space’s axes. We demonstrate that each of these ASFAs is generated from a group of functionally related genes by showing that genes associated with the same axis form more densely connected subnetworks (higher clustering coefficient) in the original human PPI network than randomly chosen genes (Mann-Whitney U test with p-value of 6.46 × 10^−28^). Furthermore, we demonstrate that these newly generated ASFAs are biologically coherent and complement the ASFAs derived from GO BP terms. Thus, we show that all axes of the embedding space are functionally relevant.

In conclusion, by analyzing the biological coherence of the ASFAs, we demonstrate that the axes of the embedding space capture coherent complex cellular functions from the functional organization of the embedding space. These results open a new opportunity for the development of data-driven ontologies using the set of ASFAs to summarize the cell’s functional organisation.

### The axes of the embedding space uncover the human evolutionary history

Having demonstrated that our ASFAs represent coherent functions of the human cells, we investigate if they can be used to get insights into humans’ evolutionary history. To this end, we propose to investigate the link between the ASFAs and evolution. We introduce the concept of “conservation degree,” which quantifies the evolutionary conservation of a given GO BP term by the number of different taxons in which it appears (detailed in Methods, section Quantifying the evolutionary conservation of biological functions). The higher the conservation degree of a GO BP, the more evolutionary conserved it is. We extend this concept to ASFAs: for a given ASFA, the conservation degree is the union of the different taxons in which the GO BP terms associated with its corresponding axis appear. We divide the ASFAs according to their conservation degree into three classes:“prokaryotes,” “eukaryotes” and “vertebrates” (detailed in Methods, section Quantifying the evolution conservation degrees of the ASFAs). We end up with 156 (53%), 101 (35%), and 31 (10%) ASFAs classified as “prokaryotes,” “eukaryotes” and “vertebrates” in the human PPI network embedding space, respectively. We analyze the meaning of these groups of ASFAs in the context of evolution.

We find that “prokaryotes” ASFAs define highly conserved functions in evolution (average conservation degree of 13.7, see Figure 3). These functions connect complex human cellular functions to ancient prokaryote ones (see Table 1). For instance, axis 144’s ASFA has a conservation degree of 13. Among the taxons that are connected to this ASFA, we find several vertebrates, including rats (taxon id: 10116), mice (taxon id: 10116), and chicken (taxon id: 9031), but also bacteria, such as *E. coli* (taxon id: 511145). This suggests that the biological function represented by this ASFA may have originated in prokaryotes, but is conserved across evolution. Indeed, it describes the regulation of neuronal synapses in vertebrates by gamma-Aminobutyric acid. Interestingly, the sets of proteins comprising synapse receptors, signaling, and biosynthetic pathways necessary for this regulation arose in prokaryotes to enable prokaryotic organisms to adapt to changing environments [38, 39].

**Fig. 3:**
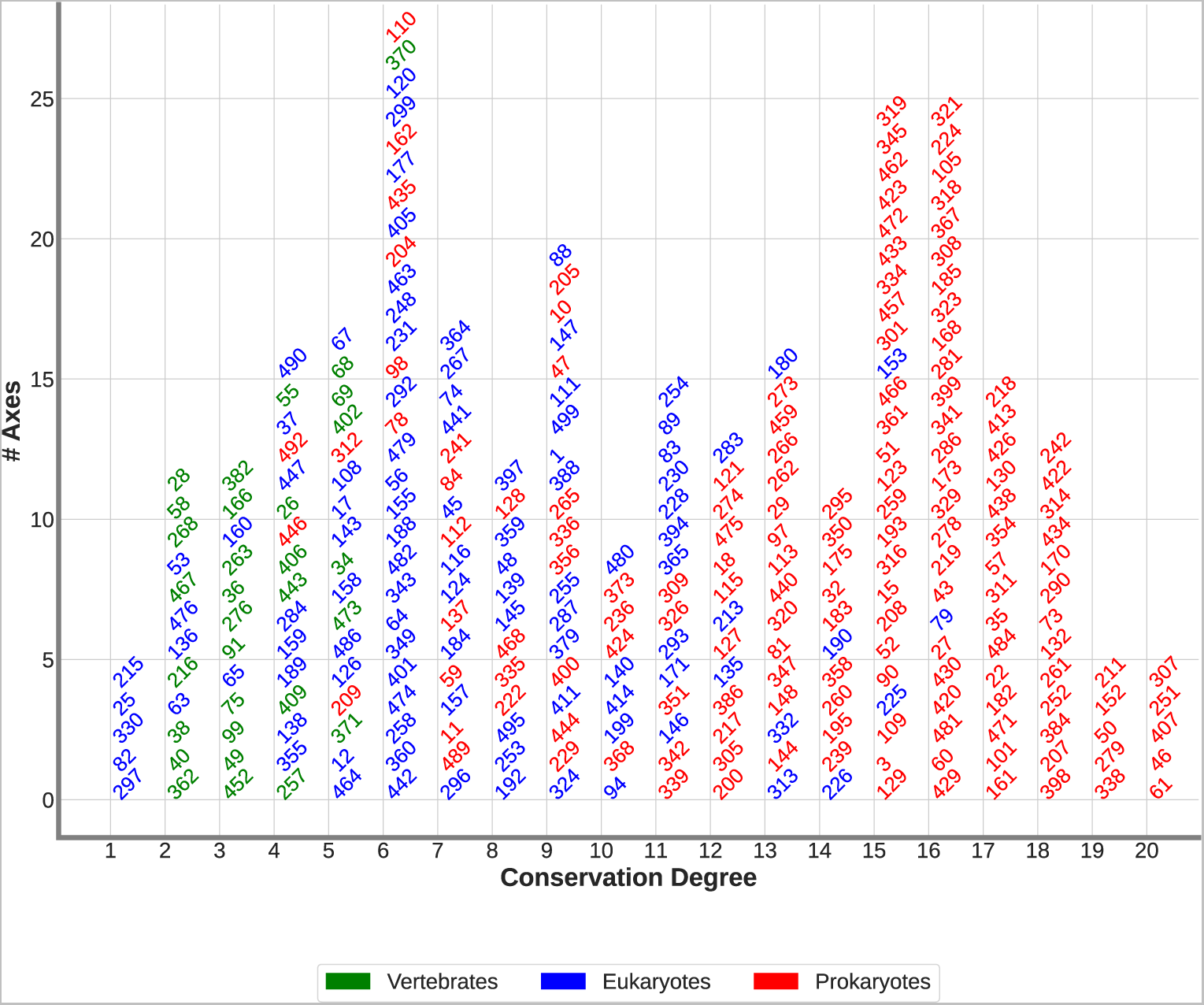
The human ASFAs give insights into the evolutionary history of humans. We use the conservation degree of the ASFAs to divide them into three groups: “prokaryotes,” “eukaryotes,” and “vertebrates” (color-coded). Then, we order the ASFAs according to their conservation degree. The horizontal axis displays the conservation degree of the ASFAs. The vertical axis shows the number of ASFAs with a certain conservation degree. Each ASFA is represented in the plot by the number of the axis from which it was obtained.

In contrast, “eukaryotes” ASFAs are newer in evolution with an average conservation degree of 7.3, which is lower than that of “prokaryotes” ASFAs (13.7, see Figure 3). These ASFAs uncover evolutionary connections between humans and other eukaryotes (see Table 1). For instance, axis 64’s ASFA sheds light on the evolutionary divergence in neurogenesis. In particular, it is connected to the embryonic stem cell differentiation into neural crest. Surprisingly, although this process is considered a functional innovation of vertebrates, we find that this ASFA is connected to two invertebrates: fruit fly (taxon id: 7227) and worm *C. elegans* (taxon id: 6239). To understand this observation, we analyze the three GO BP terms associated with axis 64 and find that they are connected to stem cell fate differentiation (GO:0001708 and GO:0048866) and neural crest stem cell differentiation (GO:0014036). Interestingly, GO:0001708 and GO:0048866 appear in fruit fly and *C. elegans*, which supports the hypothesis that the regulatory programs involved in neural crest formation evolved from programs already present in the common vertebrate-invertebrate ancestor [40]. Indeed, recently, a group of cells in invertebrates was identified with the characteristics of the neural crest ones [41].

Finally, the “vertebrate” ASFAs are on average the newest in the evolutionary history of humans (average conservation degree of 3.4, see Figure 3). In general, they describe specific traits that are unique to vertebrates. For instance, we find eight ASFAs that define functions connected to the development of tissues that are unique to vertebrates, such as cranial development, bone remodeling, skeletal muscle and cardiac muscle (see axes 68, 402, 36 and 406 in Table 1, respectively).

In conclusion, we demonstrate that each axis of the embedding space represents a well-defined function of the human cell. Moreover, by analyzing our new ASFAs, we find evolutionary connections between different species. We find similar results for other species as well (see Supplementary Results, section The Axes of the embedding space give insights into the evolutionary history of species, for more examples of human and the other species).

## Discussion

By introducing our new axes-based method, we shift the exploration of the gene embedding spaces’ organization from the genes’ embedding vectors to the axes of the embedding space. This is the first study that does not discard the axes of the gene embedding space; instead, we demonstrate that they can be used to decipher biological information from the gene embedding space. We show that our axes-based method outperforms the classic gene-centric approach in capturing the cell’s functional organization. Moreover, it also surpasses our previous FMM-based approach by enabling the identification of significant functional interactions between GO BP terms, rather than simply capturing all interactions.

Furthermore, we show that our methodology captures new interactions between pairs of GO BP terms that are not described in the Gene Ontology, but are biologically coherent. We demonstrate that the interactions captured by each axis represent a higher-order cellular function (a.k.a. ASFAs) and their combination offers a summarized functional fingerprint of the cell. This fingerprint can go from a generic overview of the cell to the most specific one depending on the number of dimensions used for generating the gene embedding space. Finally, we leverage our methodology to get insights into the evolutionary history of different species.

Due to the incompleteness of GO annotations, not all axes have GO BP terms associated, i.e., our methodology can not find their meaning. We overcome this limitation by associating genes with the embedding space’s axes and using their gene descriptions to generate the corresponding ASFAs and demonstrate that these newly generated ASFAs are biologically coherent and complement the ASFAs derived from GO BP terms.

To conclude, our methodology could be easily applied to other bioinformatics tasks, such as the development of molecular network data-driven ontology (using the ASFAs as functional annotations and connecting them based on their similarity), or as the bases for molecular network drawing algorithms by using the axes to summarize the functional organization of molecular networks. Furthermore, our methodology is generic and can be applied to any discipline that analyzes the organization of networks by using network embeddings, e.g., social, or economic networks, paving the road to new algorithms for functionally mining the data by utilizing the axes of the embedding space.

## Methods

### Biological datasets

#### PPI Networks

We collect the experimentally validated PPIs of *Homo sapiens sapiens* (human) and the PPIs of the five most frequently used model organisms: *Saccharomyces cerevisiae* (budding yeast), *Schizosaccharomyces pombe* (fission yeast), *Rattus norvegicus* (brown rat), *Drosophila melanogaster* (fruit fly) and *Mus musculus* (house mouse) from BioGRID v.4.2.191 [42]. We model these species PPI data as PPI networks in which nodes represent protein-coding genes, and edges connect nodes (genes) whose corresponding protein products physically bind. The statistics of these PPI networks are presented in Supplementary Table 1.

#### Network Representation

We represent the PPI networks with their Positive Pointwise Mutual Information (PPMI) matrices, *PPMI*. These matrices quantify how frequently any two nodes in the corresponding PPI network co-occur in a random walk compared to what would be expected if the occurrences of the nodes were independent. Following Xenos *et. al* [43] and Doria-Belenguer *et. al* [25], we use the Deepwalk closed formula by Qiu *et. al* [44] with its default settings to compute the PPMI matrix (see equation 1):

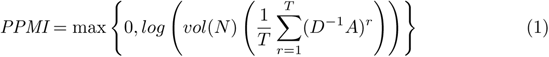

where *A* is the adjacency matrix of the network *N*, *D* is the diagonal matrix of *A*, *vol*(*N*) is the volume of *N*, *T* = 10 is the length of the random walk.

This formula can be interpreted as a diffusion process that captures higher-order proximities between the nodes in the network; hence, the PPMI matrix is a richer representation than the adjacency matrix [43]. As demonstrated by Xenos *et al.* [43] and Doria-Belenguer *et al.* [25], the extra information encoded in PPMI matrices leads to embedding spaces that better functionally organize the vectorial representation of both genes and gene functions than those generated by using the adjacency matrix.

#### Biological Annotations

We use the GO BP terms to represent the biological functions of a cell [20]. We collect the experimentally validated GO BP terms from NCBI’s FTP server (gene2go file, collected on 28 September 2021). To better capture the higher level functional organization of the cell, we not only annotate the genes with the GO BP terms that they are associated with in the gene2go file, but also with the ancestors of these terms in GO ontology. To uncover these ancestor terms, we use GOATOOLS [45] and follow the ‘is a’ and ‘part of’ links between the GO terms in the ontology’s directed acyclic graph (go-basic.obo file, collected on 04 November 2021 from the GO website). Supplementary Table 2 shows the total number of GO BP terms that annotate genes in each species PPI network. From the same gene2go file, we also keep the information about the species (taxons) in which each annotation appears after considering extension with ancestor terms (out of the 20 taxons included in the file).

### Embedding the PPI networks

To obtain a PPI network embedding space, we use three different network embedding algorithms: NMTF [15], ONMTF (detailed below) and Deepwalk [6].

#### NMTF and ONMTF

We use NMTF to decompose the PPMI matrix representation of a molecular network, *X*, as the product of three non-negative factors: *PPMI ≈ P × S × B^T^*, where rows of the matrix *E* = *P × S* define the set of embedding vectors of the genes, and the columns of *B* define the basis (a.k.a, axes) of the space in which the genes are embedded [46]. We use NMTF with and without applying the orthonormality constraint (“ONMTF” and “NMTF,” respectively), *B^T^× B* = *I* to the basis-defining matrix, *B*. This constraint leads to minimal co-linearities (hence, dependencies) between the axes of the embedding space [47]. The ONMTF and NMTF decompositions are done by minimizing functions 2 and 3, respectively:

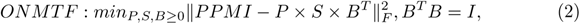

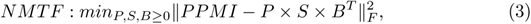

where F denotes the Frobenius norm. These optimization problems are NP-hard [15]; thus, we heuristically solve them by using a fixed point method that starts from an initial solution and iteratively uses multiplicative update rules [15]. Such rules guarantee convergence towards a locally optimal solution that verifies the Karush-Kuhn-Tucker (KKT) conditions [15] (detailed in Methods, section Fixed point method with multiplicative update rules). To generate initial *P*, *S*, and *B* matrices, we use the Singular Value Decomposition-based strategy [48]. This strategy makes the solver deterministic and reduces the number of iterations needed to achieve convergence [48]. To measure the quality of the factorization, we compute the Relative Square Error (RSE) between the input matrix, *PPMI*, and its corresponding decomposition, *PSB^T^*, as 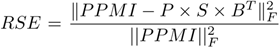. We stop the iterative solver when the value of the RSE is not decreasing anymore, or after 500 iterations.

#### Deepwalk

We use Deepwalk with its default settings [6] to learn the embedding vectors of the genes produced by this algorithm. Similar to NLP-based network embedding algorithms, Deepwalk learns these vectors by considering the node paths traversed by random walks in the PPI network as sentences and leveraging a skip-gram neural network for learning the embedding vectors of the nodes (genes) [6]. Note that Deepwalk only computes and outputs the gene embedding matrix, *E*_1_, but not the basis of the space in which the genes are embedded, *B*_1_. However, it has been demonstrated that Deepwalk, like other NLP-based network embedding algorithms, is implicitly factorizing a random walk mutual information matrix, *Y* [44]. Thus, Deepwalk approximately solves the decomposition of *Y* as 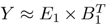. Following Qiu *et. al* [44] and Levy *et. al* [49], we approximate *Y* by its corresponding PPMI matrix by using equation 1. Then, we obtain the basis from Deepwalk embeddings by observing that in Deepwalk’s implicit decomposition: 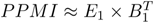, both *PPMI* and *E*_1_ are fixed, which leads to a closed formula for computing *B*_1_: 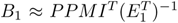, where −1 denotes the Moore-Penrose pseudoinverse. Importantly, the implicit decomposition from Deepwalk has fundamental differences from those of NMTF. First, it is not constrained to be non-negative, i.e., the coordinates of the embedding vectors of the genes, *E*, can be positive, negative, or equal to zero. Second, the basis, *B*_1_, can not be constrained to be orthonormal, so it results in more correlated axes. In other words, Deepwalk decomposition has more degrees of freedom than that of NMTF and ONMTF, which may affect the topology of the gene embedding space.

### Fixed point method with multiplicative update rules

As presented in Methods, section Embedding the PPI networks, the ONMTF and NMTF can be formulated as the minimization problem shown in Equations 2 and 3, respectively. These optimization problems are NP-hard [15], thus to solve them we use a fixed point method that starts from an initial solution and iteratively uses the following multiplicative update rules [15], derived from the Karush-Kuhn-Tucker (KKT) conditions, to converge towards a locally optimal solution:

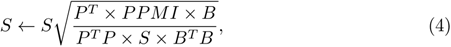

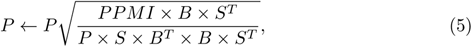

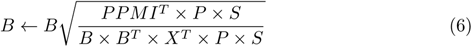

We start from initial solutions, *S_init_*, *P_init_*, *B_init_*, and iteratively use Equations 4, 5 and 6 to compute new matrix factors *S*, *P* and *B* until convergence. To generate initial *S_init_*, *P_init_* and *B_init_*, we use the Singular Value Decomposition based strategy [50]. However, SVD matrix factors can contain negative entries; thus, we use only their positive entries and replace the negative entries with 0, to account for the non-negativity constraint of the NMTF and the ONMTF. This strategy makes the solver deterministic and reduces the number of iterations needed to achieve convergence [50].

### Annotating the axes of the gene embedding space with GO BP terms

We propose to use the axes of the space in which the genes are embedded to capture the most relevant interactions between the functional annotations, which we also embed in the gene embedding space. Our method takes as input: the matrix factor, *B*, which contains the axes of the gene embedding space, and the relation-matrix between the genes and their GO BP functional annotations, *A*, in which entry *A*[*i, j*] is one if annotation *a_i_* annotate gene *g_j_*, and it is zero otherwise.

First, we generate the embedding vectors of the functional annotations in the gene embedding space. To this end, we decompose matrix *A* as the product of two matrix factors, *U* and *B^T^*, *A ≈ U × B^T^*, where rows of matrix *U* (that we call *u_i_*) are the embedding vectors of the annotations, *a_i_*, in the gene embedding space spanned by the basis, *B*, i.e., entry *u_i_*[*j*] corresponds to the coordinate of the vector *u_i_* with respect to the axis *j* in *B*. Since matrices *A* and *B*are known, we directly compute *U* by: *U ≈* (*B^T^*)^−1^ *× A*, where (*B^T^*)^−1^ is the Moore-Penrose pseudoinverse of *B^T^* [51]. Then, we associate annotation *a_i_* to axis *j* if the value of the projection of *a_i_* on *j*, *u_i_*[*j*], is statistically significantly larger than expected at random. We assess this statistical significance by performing the following bootstrapping-based method with 100,000 iterations. In each iteration, we randomly shuffle the functional annotation matrix *A* and use it as input to obtain the random matrix, *R*, which rows (that we call *r_i_*) are the random embedding vectors of the annotations, *a_i_*, in the gene embedding space spanned by the basis, *B* i.e., entry *r_i_*[*j*] corresponds to the coordinate of the vector *r_i_* with respect to the axis *j* in *B*. After all the iterations, the p-value of entry *u_i_*[*j*] is computed as 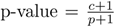 where *c* corresponds to the number of times that the observed value of *u_i_*[*j*] is lower than or equal to *r_i_*[*j*] and *p* corresponds to the number of iterations (100,000 in our analysis). For each annotation, we correct the resulting p-value for multiple hypothesis testing by applying the False Discovery Rate (FDR [52]) method over all axes. We consider the projection of annotation *a_i_* on axis *j* to be statistically significant if its corrected p-value is lower than or equal to 5%. Finally, following the hard clustering procedure introduced by Brunet *et. al* [53], we consider that annotation *a_i_* is associated with axis *j* if *u_i_*[*j*] is statistically significant and is the entry with the maximum value in vector *u_i_*.

### Quantifying the evolutionary conservation of biological functions

During evolution, cellular functions can be conserved, lost, or gained by the species (or taxons) via natural selection. To quantify the evolutionary conservation of a given biological function (represented by a GO BP annotation in this study), we introduce the “conservation degree,” which we define as the number of different taxons in which the annotation appears (out of the 20 taxons available in the gene2go file obtained from NCBI’s FTP server, detailed in Methods, section Biological datasets). Intuitively, the higher the conservation degree of a function, the more evolutionary conserved it is (from 1 to 20).

We evaluate if the conservation degree also carries information about the specificity of the function represented by a GO BP term (if it is a high-level, or a specialized cellular function) by computing the Pearson’s correlation coefficient [54] between our conservation degrees and two known measures of functional specificity: the number of genes that are annotated by a particular GO BP term (“number of genes,” for short) and the level of the GO BP terms in the GO hierarchy (“level,” for short). GO BP terms that represent generic cellular functions annotate a large number of genes and are at lower levels in the GO hierarchy. In contrast, GO BP terms that annotate a low number of genes and are at higher levels in the GO hierarchy, represent more specialized cellular functions. We find that the conservation degree is positively correlated with the number of genes (Pearson correlation coefficient of 0.44 with a p-value ≤ 1 × 10^−323^) and negatively correlated with the level (Pearson correlation coefficient of *−*0.27 with a p-value ≤ 1 × 10^−323^). Thus, a high conservation degree relates to generic functions that annotate larger sets of genes, while low conservation degrees relate to more specific functions that annotate smaller sets of genes.

Also, we investigate if the conservation degrees of the GO BP terms relate to their embedding vector positions in the embedding space. To this end, we embed GO BP terms into the species PPI network embedding spaces (detailed in Methods, sections Embedding the PPI networks and Annotating the axes of the gene embedding space with GO BP terms) and we study the correlation between the mutual positions of their embedding vectors in the embedding space (measured by their pairwise Euclidean distances) and their conservation degree. For a given GO BP term, we observe that the greater its conservation degree, the farther its vectorial representation is from the rest of the GO BP terms in the embedding space (Spearman correlation coefficient of 0.72 with p-value ≤ 1×10^−323^). Interestingly, we find that after a specific conservation degree, the average pairwise Euclidean distance drastically increases in all species PPI network embedding spaces. For instance, after reaching a conservation degree of 17 in the human PPI ONMTF embedding space, the average pairwise Euclidean distance drastically increases from 1.20 to 16.48 (see Supplementary Figure 1). Thus, we use this observation to divide the GO BP terms into three categories: “specific” (conservation degree between 17 to 20), “generic” (conservation degree between 1 to 4), and “background” (GO terms that are neither generic nor specific) for human ONMTF embedding spaces. These results hold for the other species and embedding algorithms (see Supplementary Figure 1).

### Assessing the ability of our axes-based methodology to capture biological information

Current embedding approaches rely on the organization of the genes embedding vectors in the embedding space to uncover the cell’s functional organization from molecular networks. In particular, they apply functional enrichment analysis to identify those GO BP terms that are statistically overrepresented in the clusters of genes. The GO BP terms that are statistically enriched in each cluster are then summarized to capture the cell’s functional organization. The ability of these approaches to uncover the cell’s functional organization is usually quantified by the number of gene clusters enriched in GO BP terms (“enriched clusters”), the number of GO BP terms enriched across these gene clusters (“enriched GO BP terms”), and the semantical similarity of GO BP terms enriched in the same cluster.

Instead of using the organization of the embedded entities (genes and genes’ functions) in the embedding space, in this paper, we propose using the axes of the embedding space in which the entities are embedded to uncover the cell’s functional organization from the space in which molecular networks are embedded. Similar to the gene-centric approach, we evaluate the ability of our axes-based method to capture the cell’s functional organization by analyzing the amount of GO BP terms that are associated with them. We report both: the percentage of the total GO BP terms that are associated with the axes (“% GO BP”) and the percentage of axes with at least one associated GO BP term (“% Axes”). We also investigate whether this captured biological knowledge is coherently stratified across the axes, i.e., if GO BP terms associated with the same axis are more functionally similar than those associated with different axes. To this end, we compute Lin’s semantic pairwise semantic similarity [55] between any two GO BP terms. This measure captures the similarity in the biological concepts represented by the GO BP terms, i.e., a high semantic similarity indicates that two GO BP terms are functionally related. We term “intra-SEmantic SImilarity” (intra-SeSi) the average semantic similarity of the pairs of GO BP terms that are associated with the same axis, and “inter-SEmantic SImilarity” (inter-SeSi) the average semantic similarity of the pairs of GO BP terms that are associated with different axes. We report how many times the intra-SeSi and inter-SeSi are higher than the expected random semantic similarity and the p-value of the corresponding one-tailed Mann-Whitney U test. Alternatively, we apply the same methodology, but replace Lin’s semantic similarity with the shortest path distance in the ontology DAG as a measure of functional similarity between the GO BP terms. The lower the shortest path distance between two GO BP terms, the more functionally related they are.

### Generating the Axes-Specific Functional Annotations

To obtain annotations that globally summarize the biological functions captured by each axis of the embedding space, we propose to use the GO BP terms captured by the axes to generate new data-driven functional annotations, which we call *Axes-Specific Functional Annotations* (ASFAs). These ASFAs are a set of keywords extracted from the text descriptions of the GO BP terms that are associated with a given axis that summarize them. To find these keywords, we adapt the Term Frequency Inverse Document Frequency (TF-IDF) used in the NLP field. The TF-IDF is a statistic that reports how important a word is to a document (e.g., chapters of a book) in a corpus (e.g., a textbook) [56]. We extend this statistic to our problem at study by considering the union of all the GO BP terms text for the GO BP terms that are associated in all axes of the embedding space as the corpus. On the other hand, we consider as a document of this corpus the union of the text descriptions of the GO BP terms that are associated with an axis (ideally having as many documents as axes). Then, we compute the TF-IDF of a word in a document by applying equation 7:

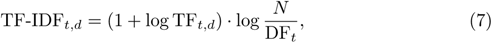

where t is a word in the document, d, *TF − IDF_t,d_*is the number of occurrences of t in d, *DF_t_* is the number of documents containing t, and N is the total number of documents.

Since not all the words add semantic meaning to a text (e.g., “where” or “of”), they could add noise to the TF-IDF, i.e., these so-called “stop words” are usually removed before applying the TF-IDF [57]. Stop words that we consider are all those that are in the list of stop words from the NLTK package version 3.6.7 [58]. Similarly, we filter all the “stop words” from the text definitions of the GO BP terms before computing the TF-IDF. Finally, for each axis, we build its ASFA by taking all the words with a TF-IDF higher than 0. This is because a word with a TF-IDF higher than 0 is considered to have a level of significance for the document. On the other hand, a word with TF-IDF equal to 0 indicates that the word is not considered important for the document based on the collection of documents.

Since GO ontology is incomplete, many genes lack GO BP term annotation. For instance, only 48.6% of the genes in the human PPI network are annotated with GO BP terms. Because of this, many genes are left unannotated (with no associated GO terms). Thus, some axes may not capture any embedded functional annotations from the embedding space. To overcome this issue, we propose to associate genes with the embedding space’s axes and use the text descriptions of the genes from the Alliance of Genome Resources database v.5.2.1 to generate the corresponding ASFAs. To this end, we associate each gene to the axis for which the projection of the gene’s embedding vector has the largest value (in the spirit of the hard clustering procedure of [53]). Finally, having a set of gene descriptions for each axis, we apply the same TF-IDF-based approach used with the GO BP functional annotations to build their ASFAs. In particular, we consider all the genes associated with the axes of the embedding space (all their gene descriptions) as the corpus. On the other hand, we consider as a document of this corpus the union of the gene descriptions of the genes that are associated with an axis (ideally having as many documents as axes). Then, we compute the TF-IDF of a word in a document by applying the equation 7.

### Quantifying the evolution conservation degrees of the ASFAs

To gain insights into the human evolutionary history, we propose to investigate the link between the ASFAs and evolution. To this end, we extend the notion of the conservation degree of a GO BP term (defined in Methods, section Quantifying the evolutionary conservation of biological functions) to an ASFA. For an individual ASFA, we obtain its conservation degree by the union of the different taxons in which the GO BP terms associated with its corresponding axis appear. Also, we search for evolutionary patterns across our ASFAs by identifying those ASFAs that describe biological functions that are conserved from prokaryotic organisms, that appeared for the first time in eukaryotes, or that are unique to vertebrates. To this end, for a given ASFA, we take the union of the different taxons in which the GO BP terms associated with its corresponding axis appear. Then, we perform a systematic classification of these taxons into two life domains: *Bacteria*, which represents prokaryotic organisms, or *Eukarya*, which represents eukaryotic organisms. For taxons that belong to the *Eukarya* domain, we also identify those taxons that are part of the sub-phylum *Vertebrata*. Based on the systematic classification of its taxons, we consider the ASFA to be related to “prokaryotes” if at least one of the taxons is a prokaryote, “eukaryotes” if all the taxons are eukaryotes, but none of them is vertebrate, and “vertebrates” if all the taxons are eukaryotes, but at least one of them is a vertebrate. Note that the intuition behind this classification is to find from which organisms (prokaryotes, eukaryotes, or vertebrates) the functions represented by the ASFAs of a given species were inherited (e.g., the human ASFAs).

## Data and code availability

Data and source code can be accessed at https://gitlab.bsc.es/sdoria/axes-of-biology.git

## Acknowledgments

This work was supported by the European Research Council (ERC) Consolidator Grant 770827 and the Spanish State Research Agency AEI 10.13039/501100011033 grant number PID2019-105500GB-I00.

## Declarations

The authors declare no competing interests.

