## Supplementary Data for "The axes of biology: a novel axes-based network embedding paradigm to decipher the functional mechanisms of the cell"

#### Supplementary Results

##### Orthonormality and positive constraints improve the functional organization of the gene embedding space

In Results, section The axes of the embedding spaces capture the cell's functional organization of the main manuscript, we demonstrated that the axes of the ONMTF embedding spaces capture more and better-stratified functional information than the other methods. Here, we analyze if the ability of ONMTF to produce embedding spaces whose axes capture more, and better stratified functional information can be attributed to the properties of the embedding spaces produced by the ONMTF. ONMTF embedding spaces have two properties, orthonormality, and non-negativity. We assess the effect of these properties in disentangling functional knowledge from the biological networks. Since the embedding space is orthonormal, its axes should represent non-ambiguous and non-dependant directions of the space. We confirm this first property by computing the average pairwise cosine similarity in-between the axes of the ONMTF, NMTF, and Deepwalk embedding spaces. It is important to note that Deepwalk embedding spaces are not constrained to be positive, which means that the cosine similarity is bounded from -1 to 1 instead of from 0 to 1. Thus, to make it comparable to the NMTF and ONMTF spaces, we report the absolute pairwise cosine similarity in-between their axes. A cosine similarity of 1 indicates that two axes are identical (i.e., redundant), and a value of 0 indicates that the axes are orthogonal (i.e., perpendicular).

We observe that the axes of the NMTF embedding spaces have, on average, the largest number of similar axes (average pairwise cosine similarity of 0.986), followed by Deepwalk (average pairwise cosine similarity of 0.90), and ONMTF (average pairwise cosine similarity of 0.24). These results suggest that the majority of the axes in the NMTF embedding space are redundant, i.e., some dimensions do not contribute to disentangling functional knowledge from the biological networks. This high redundancy, in turn, explains the low percentage of GO BP terms associated with the axes of NMTF spaces (11.95%) in comparison to ONMTF (37.12%). Also, we see that, although the axes of the Deepwalk spaces are not constrained to be orthonormal, their axes have a lower average pairwise cosine similarity (average of 0.10) than the ones of the NMTF. We explain this observation by the degrees of freedom of Deepwalk spaces. In other words, since Deepwalk spaces are not constrained to be positive, the chance that two random vectors are identical is lower than in NMTF spaces. Finally, we observe the absence of non-negativity constraints in Deepwalk embedding spaces decreases its ability to capture the cell's functional organization (GO BP terms less coherently stratified than ONMTF and NMTF, results presented in the previous section). We hypothesize that this observation is connected with the fact that biological processes are often non-negative and additive [1], i.e., positive embedding spaces are more suitable to capture these complex biological mechanisms.

In conclusion, the embedding in positive and orthonormal spaces, which only NMTF-based frameworks allow for, leads to the embedding spaces that best capture the cell's functional organization from the biological networks.

#### Exploring the impact of the network embedding space's dimensionality on the biological information captured by the axes

In Results, section The axes of the embedding spaces capture the cell's functional organization of the main manuscript, we demonstrated that the embedding axes capture functional knowledge from network embeddings (represented by GO BP terms). In this section, we investigate how the space's dimensionality affects the specificity of the GO BP terms captured by the axes, the amount of GO BP terms captured by the axes, the number of axes with at least one associated GO BP term and the coherence of the stratification of the GO BP terms across the axes. In particular, to analyze the impact of the dimensionality on the specificity of the GO BP terms captured by the axes, we divide them into three groups: "specific," "generic," and "background" (detailed in Methods, section Quantifying the evolutionary conservation of biological functions). We measure how well these "specific," "generic," and "background" GO BP terms are captured by the axes as a function of the dimensionality of the embedding space (fold increase with respect to the lowest dimensional space with 50 dimensions).

We find that most of the "generic" functions (average of 90%) are associated with the axes of human lowest dimensional embedding space (50 dimensions). Importantly, we find that increasing the dimensionality of the embedding space allows us to capture more "generic" functions (fold increase remains close to 1, see Supplementary Figure 2). In contrast, increasing this dimensionality allows for capturing more "background" and "specific" functions, with the specific ones benefiting the most from the increase in the number of dimensions (see Supplementary Figure 2). Moreover, increasing the dimensionality enhances the stratification of biological information captured by the axes, with more semantic similar GO BP terms associated with the same axis (see Supplementary Figure 3). These results suggest that the embedding space needs more dimensions to disentangle "specific" biological functions encoded in the species PPI networks.

Nevertheless, this disentanglement has a limit since after 500 dimensions there is no significant benefit in increasing the space's dimensionality. First, the number of axes capturing at least one GO BP term reduces to less than 50% and the total amount of GO BP terms captured flattens after 500 dimensions (see Supplementary Figure 1). Second, the fold increase of "specific" functions is significantly reduced after 500 dimensions (see Supplementary Figure 2). Third, the semantic similarity of GO BP terms associated with the same axis flatters after 400-500 dimensions. Thus, adding more dimensions does not improve the capture of either more biological information or more specific information from the embedding space. Interestingly, these observations are in line with the results reported in other artificial intelligence fields, such as NLP, where a low dimensionality of the word embedding fails to capture all possible word relations ("specific" relations), and after a certain number of dimensions, the embeddings can not disentangle more word relations [2]. We find similar results for the rest of the studied species ONMTF embedding spaces (see Supplementary Figures 4 and 5).

Based on these results, we consider the optimal dimensionality of a given specie-specific PPI network embedding space as the one that finds a balance between the three observations introduced above (i.e., amount of information captured, specificity of this information, and the coherence in the stratification of the information captured across the axes). Based on these criteria, we choose 500 dimensions as the optimal dimensionality for the human ONMTF embedding space (the optimal number of dimensions for the rest of species ONMTF embedding spaces can be found in Supplementary Table 6). This optimal dimensionality is coherent with the number of dimensions usually applied in NLP [3, 4].

#### Our axes-based method outperforms the classic gene-centric approach in capturing the cell's functional organization

In this section, we compare the ability of our axes-based method to uncover the cell's functional organization from PPI network embedding spaces to that of the standard gene-centric approach. To this end, we consider the six species PPI networks described in Results, section Biological datasets of the main text, which we embed by applying ONMTF, NMTF, and Deepwalk embedding algorithms (as detailed in Methods, section Embedding the PPI networks of the main text). We generate these embedding spaces with increasing dimensionalities (from 50 to 1000 dimensions with a step of 50).

In a first step, we apply the standard gene-centric approach to uncover the cell's functional organization from the species PPI network embedding spaces described above. To this end, we perform the following gene clustering and enrichment analysis. For each embedding space, we cluster together genes that are embedded close in space by applying the k-medoid algorithm [5] on the pairwise cosine distances of the genes' embedding vectors. For the number of clusters, we use the heuristic rule of thumb ($k = \sqrt{\frac{n}{2}}$, where $n$ is the number of nodes in the species network) [6]. We end up with 95, 54, 40, 67, 63 and 38 clusters for human, budding yeast, fission yeast, fruit fly, mouse and rat, respectively. Then, we measure the enrichment of the resulting gene clusters in GO BP terms by using the sampling without replacement strategy (hypergeometric test), and we consider a GO BP term to be significantly enriched in a gene cluster if the corresponding enrichment p-value, after Benjamini and Hochberg correction for multiple hypothesis testing [7], is smaller than, or equal to 5%. As standardly done in literature, we quantify the ability of the gene-centric approach to uncover the cell's functional organization by analyzing the number of different GO BP terms that are enriched in the gene clusters, the number of clusters with at least one enriched GO BP term and the average pairwise Lin's semantic similarity of GO BP terms enriched in the same gene cluster (detailed in Methods, section Assessing the ability of our axes-based methodology to capture biological information of the main manuscript).

In a second step, we apply our axes-based methodology to uncover the cell's functional organization from the species PPI network embedding spaces described above. To this end, for each embedding space, we embed GO BP terms into the space and associate them to the axes of the embedding space (detailed in Methods, section Annotating the axes of the gene embedding space with GO BP terms). Similar to the gene-centric approach, we quantify the ability of our axes-based method to capture the cell's functional organization by analyzing the number of different GO BP terms that are associated with the embedding axes, the number of embedding axes with at least one associated GO BP term and the average pairwise Lin's semantic similarity of GO BP terms associated with the same axis (detailed in Methods, Assessing the ability of our axes-based methodology to capture biological information of the main manuscript).

To contrast the ability of our axes-based method to uncover the cell's functional organization from species PPI network embedding spaces to that of the standard gene-centric approach, we compare (1) the number of different GO BP terms that are enriched across the gene clusters to the amount of different GO BP terms that are associated to the embedding axes, (2) the number of gene clusters that have at least one GO BP term enriched to the number of embedding axes with at least one associated GO BP term and (3) the average pairwise Lin's semantic similarity of GO BP terms enriched in the same gene cluster to the average pairwise Lin's semantic similarity of GO BP terms associated in the same embedding axis.

As shown in Supplementary Table 12, over all species PPI networks, we find that, on average, 1.79, 1.10, and 1.90 times more GO BP terms associated with the embedding axes than enriched across the gene clusters for ONMTF, NMTF, and Deepwalk embedding spaces, respectively (their percentages of GO BP terms captured by each methodology are presented in Supplementary Tables 3 and 10). Thus, our axes-based methodology disentangles more biological information from the embedding space than the standard gene-centric approach. On the other hand, we do not find differences in the number between the number of axes with at least one associated GO BP term and the number of gene clusters with at least one enriched GO BP term. However, we find that our axes-based methodology not only captures more GO BP terms, but also that the functions that are associated with the same axis are functionally more coherent than the functions that are enriched in the same gene cluster (1.42, 1.10 and 1.16 times higher average semantic similarity for ONMTF, NMTF, and Deepwalk embedding spaces, respectively, see Supplementary Table 12). Hence, our axes-based methodology better captures the cell's functional organization from PPI network embedding spaces than the standard gene-centric approach.

Also, we evaluate the agreement in the biological information captured by our axes-based methodology to that by the standard gene-centric approach. To this end, we take the GO BP terms captured by both methods, i.e., the intersection between GO BP terms associated with the embedding axes and the GO BP terms enriched across the gene clusters. Then, we compare the clustering of these GO BP terms based on our axes-based methodology with that based on the standard gene-centric approach. In particular, for our axes-based method, we consider GO BP terms to cluster if they are associated with the same axis. Similarly, for the standard gene-centric approach, we consider GO BP terms to cluster if they are enriched in the same gene cluster. We measure the agreement between these two clusterings by computing the adjusted Rand Index [8]. We report the adjusted Rand Index score (see this score in Supplementary Table 13). This score is bounded between 0 and 1, where 0 corresponds to totally different clusterings and 1 to exactly similar clustering.

We find that GO BP terms associated with the same axis are not likely to be enriched in the same gene cluster (average adjusted Rand Index score of 0.14, 0.08, and 0.04 for ONMTF, NMTF, and Deepwalk embedding spaces, respectively). Thus, the axes of the embedding space capture different functional information than the gene clusters. Having observed that each method produces different clusterings of the GO BP terms, we investigate which of these clusterings provides a more coherent organization of the terms. To accomplish this, we compare the Lin's semantic similarity of GO BP terms associated with the same axis with that of those terms enriched in the same gene cluster. We find that GO BP terms associated with the same axis in the ONMTF, NMTF, and Deepwalk embedding spaces exhibit higher functional similarity (1.7, 1.13, and 1.25 times larger semantic similarity) than those enriched in the same gene cluster. Hence, our axes-based method stratifies the biological information more coherently than the standard gene-centric approach.

In conclusion, we demonstrate that our axes-based method captures more and different biological information than the standard gene-centric approach from network embeddings. Moreover, this information is more functionally coherent than the information captured by the standard gene-centric approach. Thus, our methodology outperforms the standard gene-centric approach in capturing the cell's functional organization from network embedding spaces.

#### Our axes-based method is in agreement with the FMM-based methodology

In this section, we assess if the functional interactions between the GO BP terms that are uncovered by our new axes-based methodology are supported by our previous FMM-based methodology. To this end, we consider the six species PPI networks described in Methods, section Biological datasets of the main text, which we embed by applying ONMTF, NMTF, and Deepwalk embedding algorithms (detailed in Methods, section Embedding the PPI networks of the main text). We generate these embedding spaces with increasing dimensionalities (from 50 to 1000 dimensions with a step of 50).

For a given species PPI network embedding space, our previous FMM directly quantifies all the functional interactions between any two GO BP terms that annotate genes in the PPI network by measuring the cosine distance between the GO BP terms' embedding vectors [9]. In contrast, our new axes-based only captures the significant functional interactions between GO BP terms by associating GO BP terms to the axes of the embedding space (detailed in Methods, section Annotating the axes of the gene embedding space with GO BP terms of the main text). Pairs of GO BP terms that are associated with the same axis are considered to functionally interact. For the GO BP terms that are associated with at least one embedding axis, we measure the agreement between the functional interactions uncovered by the FMM and the functional interactions that are captured by our axes-based methodology by using the following ROC curve analysis. For each GO BP pair, we consider the result of our axes-centric approach as the ground truth, i.e., a pair of GO BP terms is considered as "true" if the two terms are associated with the same axis, or as "false" otherwise. Also, for each GO BP pair, we consider as the prediction score their cosine similarity in the embedding space (1 minus their associated value in the FMM). Then, we compute the area under the ROC curve (AUROC) [10] between the ground truth and the prediction score over all the considered GO BP pairs. Note that an AUROC score of 0.5 corresponds to a random classification and a score of 1 to a perfect one. Hence, the closer to one the AUROC score, the higher the agreement between our axes-centric method and our previous FMM-based approach.

Over all species PPI networks, we find that the functional interactions uncovered by our previous FMM methodology and our new axes-based approach are in significant agreement, with an average AUROC of 0.90, 0.90 and 0.91 and all p-values $\leq 1 \times 10^{-323}$ for ONMTF, NMTF, and Deepwalk embedding spaces, respectively (see Supplementary Table 14). These results confirm that the GO BP terms that are associated with the same axis tend to be located close in the embedding space and thus, tend to have small association values in the FMM.

In conclusion, our axes-based method surpasses our previous FMM-based approach by enabling the identification of significant functional interactions between GO BP terms, rather than simply capturing all interactions. Hence, our axes-based methodology offers a more refined approach to exploring the cell's functional organization from a functional perspective.

#### Non-Annotated Axes also capture the functional mechanisms of the cell

In this section, we investigate the biological meaning of those axes without any associated GO BP term (a.k.a. empty axes). To this end, we recall that genes that form densely connected regions of a PPI network tend to share biological functions [11]. Hence, we investigate if the genes that are associated with the empty axes tend to form such densely connected neighborhoods in the human PPI network. We do this by associating genes to the 206 (41.2%) empty axes of the ONMTF human embedding space. We associate each gene to the axis for which the projection of the gene's embedding vector has the largest value (detailed in Methods, section Generating the Axes-Specific Functional Annotations of the main manuscript). Then, we evaluate the connectivity in the original human PPI network of the genes associated with the same axis by computing their clustering coefficient.

We see that the average clustering coefficient of those genes associated with the same non-empty axis (axes with associated GO BP terms) is statistically significantly higher than those genes associated with the same empty axis (Mann-Whitney U test p-value of $1.76 \times 10^{-63}$). However, we find that the average clustering coefficient of those genes associated with the same empty axis is statistically significantly higher than expected by random (Mann-Whitney U test p-value of $6.46 \times 10^{-28}$), i.e., they form more densely connected subnetworks than randomly chosen genes, which suggests that they are indeed functionally related. Hence, we explain the absence of associated GO terms on these empty axes by the lack of biological functional information (only 48.6% of the human genes in the PPI network are annotated with GO BP terms). Indeed, we find that only 40.0% of the genes that are associated with empty axes in human are annotated with at least one GO BP term. In contrast, more than half of the genes associated with non-empty axes are annotated with GO BP terms (53.8%). In other words, the empty axes capture parts of the human PPI network that have not been yet annotated. We find similar results for the rest of the studied species (see Supplementary Table 8).

To find the biological meaning of empty axes, we propose to generate their ASFAs from the text description of their associated genes rather than from the text description of their associated GO BP terms (detailed in Methods, section Generating the Axes-Specific Functional Annotations of the main manuscript). Using this approach, we obtain the ASFAs for 97.8% of the axes. We find that the interpretation of these ASFAs is less intuitive (average of 55.47 words) than the ones built using GO BP terms (average of 17.27 words), but are equally coherent. For instance, the ASFA of the empty-axis 9 is connected with the regulation of neural activity (see Table 9). Indeed, among the words that define this ASFA, we find glycine (an inhibitory neurotransmitter [12]), choline (regulator of neurological development [13]), and adenylate cyclases (regulator of the energy balance in different parts of the brain [12]). Another example is the ASFA of the empty-axis 76, which is connected to the functions of the thymus (see Table 9). This ASFA supports the observation that lipid metabolism ("chylomicron") affects lymphocyte differentiation and survival in the thymus [14].

Finally, we investigate if the ASFAs generated using genes' descriptions (a.k.a, genes' perspective) agree with those generated using functional annotations (GO terms' perspective). Interestingly, we find that the gene perspective ASFAs are not only in agreement with the GO terms perspective ones, but also complement them. For instance, from the GO terms perspective, the ASFA of axis 68 is connected to cranial development (see Table 1). In this case, the genes' perspective not only agrees with it, but also indicates that the ASFA is linked to neural tube development (see Table 9). Similarly, the genes' perspective ASFA of axis 370 complements its GO terms' perspective ASFA. From the GO terms' perspective, this ASFA is connected to the activation of natural killer lymphocytes (see Tables 1). The gene' perspective hallmarks the importance of the "glutaminyl-tRNAGln" and amidotransferase for the correct functioning of their mitochondria, which is connected to the activation of these lymphocytes [15] (see Table 9).

In conclusion, we demonstrate that all the axes of the embedding space have a coherent biological meaning. For those axes that do not have any GO BP term associated, we propose an approach method that finds the meaning of empty axes. We demonstrate that the ASFAs generated by using it agree with and complement the ones obtained by using the GO BP terms.

#### The axes of the embedding space uncover new functional interactions between GO BP terms in different model organisms

In section Results, section The axes of the embedding space uncover new functional interactions between GO BP terms of the main manuscript, we demonstrate that our axes-based methodology captures new functional interactions between GO BP terms and demonstrate that these interactions are biologically coherent by performing literature curation. To this end, we compute Lin's semantic similarity between any two GO BP terms (detailed in Methods, section Quantifying the evolutionary conservation of biological functions of the main manuscript). Then, for each axis, we take the average semantic similarity among its pairs of associated GO BP terms ("intra-axis SeSi"). By taking all the "intra-axis SeSi" over all the embedding axes, we define the distribution of "intra-axis SeSi" (see the distribution in Supplementary Figure 6). We consider an axis to have a significantly low "intra-axis SeSi" if its "intra-axis SeSi" is smaller than, or equal to the $5^{th}$ percentile of "intra-axis SeSi" distribution (see the distribution in Supplementary Figure 6). Based on this criterion, we find 13 (average "intra-axis SeSi" of 0.11), 8 (average "intra-axis SeSi" of 0.18), 6 (average "intra-axis SeSi" of 0.11), 7 (average "intra-axis SeSi" of 0.13), 12 (average "intra-axis SeSi" of 0.07), and 5 (average "intra-axis SeSi" of 0.17) axes in human, budding yeast, fission yeast, fly, mouse, and rat PPI network embedding spaces, respectively. For each of these axes, we evaluate if the interaction between its associated GO BP terms is biologically coherent. In the same Results section, we focus on the human PPI network embedding spaces and discuss the functional interactions between GO BP terms captured by three axes with significantly low "intra-axis SeSi" (axes 37, 59, and 119). In this section, we first continue this discussion for the rest of the axes with a significantly low "intra-axis SeSi" in human. Then, we extend this analysis to the embedding axes of the budding yeast PPI network embedding space. We choose human and budding yeast since their PPI networks are the most complete among the six studied species and also have the highest number of GO BP annotations (see Supplementary Tables 1 and 2).

**Analysis of the functional interactions captured by the axes of the human PPI network embedding space**:

Recall that among the 13 axes with a significantly low "intra-axis SeSi" in the human PPI network embedding space, we find 7 axes (53.8%) with functional interactions that are known to occur in humans, 3 axes (23.1%) that capture functional interactions that are described in model organisms, but not yet in humans, and 3 axes (23.1%) that capture functional interactions that are not described in the literature, but are biologically coherent.

Axes that capture functional interactions that are known to occur in humans include axes 37, 143, 11, 253, 351, 368, and 492. Globally, although these functional interactions captured by these axes are not connected based on the Gene Ontology (low semantical similarity), they are functionally coherent and describe higher-order processes that are known to occur in humans. Axis 143 has two associated GO BP terms: GO:0051482 (positive regulation of cytosolic calcium ion concentration involved in phospholipase C (PLC)-activating G protein-coupled signaling pathway) and GO:0052746 (the process of introducing one, or more phosphate groups into inositol). These GO BP terms are not connected based on the Gene Ontology (semantic similarity of 0.08), however, their functional interaction is biologically coherent since PLC and inositol are known to collaborate in the signal transduction of human cells [16].

Regarding, axis 11 has three GO BP terms associated: GO:1901837 (regulation of transcription of nucleolar large rRNA by RNA polymerase I), GO:0000027 (the ribosomal large subunit assembly), and GO:1902570 (protein localization to the nucleus) with an average semantic similarity of 0.085. RNA polymerase I is composed of multiple protein sub-units that are transcribed in the cytoplasm [17]. These proteins are imported into the nucleus, where they assemble into the RNA polymerase I complex. Once in the nucleus, the complex RNA polymerase I transcribe rRNA genes, which include the large ribosomal sub-unit [17]. Large ribosomal sub-units are known to be essential for the production of ribosomes. Thus, these three functions interact in the regulation of the transcriptional activity of RNA polymerase I by limiting its localization to the nucleus. This control is known to be key for ribosome production and assembly [18].

Axis 253, which has two associated GO BP terms: GO:0033566 (gamma-tubulin complex localization) and GO:0007020 (microtubule nucleation) with a semantic similarity of 0.09. We find that the interaction between these terms is biologically coherent since the gamma-tubulin complex is known to participate in the microtubule nucleation [19, 20]. With respect to axis 351 also has two associated GO BP terms: GO:0042136 (neurotransmitter biosynthetic process) and GO:0017004 (cytochrome complex assembly). While these functions are not connected based on the Gene Ontology (semantic similarity of 0.1), several studies state that cytochrome P450 is involved in the synthesis of neurotransmitters, such as dopamine and serotonin in the brain [21, 22].

Another axis 368 that capture functional interactions that are known to occur in humans, which has two associated GO BP terms: GO:0070914 (UV-damage excision repair) and GO:0042254 (ribosome biogenesis), with a semantic similarity of 0.1. The functional interaction between these functions is coherent since the repair of DNA lesions on ribosomal DNAs after UV irradiation is of fundamental importance for the cell to maintain ribosome biogenesis [23].

Finally, axis 492 has two GO BP terms associated: GO:0045292 (mRNA cis splicing, via spliceosome) and GO:0036245 (cellular response to menadione), with a semantic similarity of 0.09. Despite their low semantical similarity, we find the connection between these two terms to be functionally coherent since the mRNA splicing is known to be regulated by micronutrients, including menadione [24].

Axes that capture functional interactions that are described in model organisms, but not yet in humans include axes 59, 90, and 463. Axis 90 has eight GO BP terms associated that can be grouped into four clusters based on their Lin's semantical similarity. The first group (semantic similarity of 0.91) includes GO:0048660 (regulation of smooth muscle cell proliferation) and GO:1904707 (positive regulation of vascular-associated smooth muscle cell proliferation). The second group (semantic similarity of 0.85) includes GO:0030198 (extracellular matrix organization) and GO:0030199 (collagen fibril organization). The third group (average pairwise semantic similarity of 0.44) includes GO:00602702 (GINS complex), GO:0030154 (cell differentiation), and GO:0035987 (endodermal cell differentiation), i.e., is connected with the cell differentiation process. Finally, the last cluster only includes GO:001785 (peptidyl-lysine hydroxylation) which has an average pairwise semantic similarity of 0.09 with the rest of the terms associated with the axis. While these four groups of GO BP terms are not interconnected based on the Gene Ontology, we find that their interaction is functionally coherent. Vascular smooth muscle cells (SMCs) provide contractile function and structural support to blood vessels. Unlike other tissues, vascular SMCs are not terminally differentiated and display remarkable phenotypic plasticity [25]. It has been observed that the differentiation of vascular SMCs is highly influenced by the composition of extracellular matrix (ECM) [26]. However, the molecular mechanisms underlying the interaction between vascular SMCs and ECM are not yet understood in humans. Lysyl oxidase (LOX) is a key ECM-remodeling enzyme required for the hydroxylation of specific lysine residues (peptidyl-lysine hydroxylation) in collagen type I fibers [27]. Recent studies in mice showed that LOX is responsible for the proliferation and migration of the aortic vascular SMCs [28]. Hence, our results suggest that similar molecular mechanisms may regulate vascular SMC proliferation in humans.

Finally, axis 463 has three associated GO BP terms: GO:0045494 (photoreceptor cell maintenance), GO:1904970 (brush border assembly), and GO:1904106 (protein localization to the microvillus membrane). Although these three functions are not connected in the current Gene Ontology (average pairwise semantic similarity of 0.04), we find their interaction to be biologically coherent. In particular, the retinal pigment epithelium (RPE) performs highly specialized functions essential for the homeostasis of the neural retina, including photoreceptor maintenance, in different model organisms [29, 30]. All these homeostasis functions involve the RPE apical microvilli. The last axis, axis 59 is discussed in the main manuscript.

To conclude, axes that capture functional interactions that are not described in the literature, but are biologically coherent include axis 116, 441, and 222. Axis 441 has two associated GO BP terms: GO:0008589 (regulation of smoothened signaling pathway) and GO:0044375 (regulation of peroxisome size). These two GO BP terms are not connected based on the current Gene Ontology (semantic similarity of 0.13), however, we find their interaction to be functionally coherent. In particular, smoothened is a protein that participates in the hedgehog signaling pathway [31]. Studies in human mesenchymal stem cells revealed that hedgehog interferes with adipocyte differentiation by targeting the peroxisome proliferator-activated receptor (PPAR) [32]. Moreover, studies in the liver also hallmarked the importance of the hedgehog in controlling the peroxisomal fatty acid $\beta$-oxidation rate [33]. Thus, we hypothesize that the hedgehog signaling pathway could have an important role in lipid metabolism by regulating peroxisomes.

Regarding, axis 222 has thirteen GO BP terms associated that can be grouped into seven clusters based on their Lin's semantical similarity. The first group (semantic similarity of 0.96) includes GO:0097252 (oligodendrocyte apoptotic process) and GO:0034349 (glial cell apoptotic process). Since oligodendrocyte is part of the glial cells, this group describe the apoptosis in such cells. The second group (average pairwise semantic similarity of 0.45) includes GO:0043456 (pentose-phosphate shunt), GO:1905856 (negative regulation of pentose-phosphate shunt), GO:1990248 (regulation of transcription from RNA polymerase II promoter in response to DNA damage), and GO:0045899 (positive regulation of RNA polymerase II transcription preinitiation complex assembly). The pentose-phosphate pathway is required for the synthesis of nucleotides that are needed for transcription [34], i.e., these GO BP terms are connected to the molecular mechanisms that regulate gene expression in response to DNA damage. The third group (semantic similarity of 0.96) clusters GO:0071480 (cellular response to gamma radiation) and GO:0010332 (response to gamma radiation). The fourth group (semantic similarity of 0.97) includes GO:0072717 (response to actinomycin D) and GO:007271 (response to actinomycin D). Actinomycin D is a well-known drug that induces apoptosis and inhibits the growth of cancer cells [35]. The fifth group (semantic similarity of 0.97) includes GO:0090403 (oxidative stress-induced premature senescence) and GO:0090400 (stress-induced premature senescence). So far, the five groups of GO BP terms can be easily connected since their functional interaction could represent the induction of apoptosis in response to DNA damage in glial cells. However, apart from these five groups, we also find other associated GO BP terms that can not be easily connected with non of them. In particular, GO:0048539 describes bone marrow development. We hypothesize the connection of these GO BP terms could be attributed to the bone marrow transdifferentiation to neural cells [36]. After the apoptosis of glial cells, the differentiation of bone marrow cells into glial cells could assist in the repair of the tissue.

**Analysis of the functional interactions captured by the axes of the budding yeast PPI network embedding space**:

We find 8 axes with a significantly low "intra-axis SeSi" in the budding yeast PPI network embedding space. Among them, 6 axes (75%) have functional interactions that are known to occur in budding yeast and 2 (axes (25%) capture functional interactions that are not described in the literature, but are biologically coherent.

Axes that capture functional interactions that are known to occur in budding yeast include axes 31, 66, 107, 125, 155 and 174. Axis 31 has three associated GO BO terms: GO:0010499 (proteasomal ubiquitin-independent protein catabolic process), GO:0051131 (chaperone-mediated protein complex assembly), and GO:0080129 (proteasome core complex assembly). Although these three terms have an average semantic similarity of 0.22, their interaction is functionally coherent since they describe the proteasome core complex assembly by chaperones and the functions of this complex.

Regarding, axis 107 has two associated GO BP terms: GO:0006998 (nuclear envelope organization) and GO:0055088 ( lipid homeostasis). These terms are semantically dissimilar (semantic similarity of 0.06). However, the link between nuclear lipid homeostasis and the nuclear envelope organization has been widely investigated in different organisms. For instance, a study in human's liver demonstrated that specific subdomains of the nuclear envelope are involved in nuclear lipid homeostasis [37]. This observation has been also reported in different organisms, including budding yeast [38, 39].

With respect to axis 125, it has seven associated GO BP terms. By analyzing these GO BP terms, we find that the axis represents different cellular defense responses to methylmercury. In particular, GO:0051597 (response to methylmercury) and GO:0071406 (cellular response to methylmercury) are connected with the cellular response to methylmercury (semantic similarity of 0.97). Methylmercury is an extremely toxic organometallic cation that interferes with cell cycle progression by disrupting the organization of microtubules. This organization is represented by GO:0051382 (kinetochore assembly) and GO:0051383 (kinetochore organization). Recent studies have reported that the ubiquitin-proteasome system is involved in defense against this toxic element. This ubiquitin-proteasome response is represented by GO:0031146 (SCF-dependent proteasomal ubiquitin-dependent protein catabolic process), GO:0045116 (protein neddylation), and GO:0000338 (protein deneddylation) [40].

Regarding axis 155, it has two associated GO BP terms: GO:0006457 (protein folding) and GO:0007021 (tubulin complex assembly). Despite their low semantic similarity (0.09), their interaction is functionally coherent since it represents the folding and assembly of the tubulin complex.

As for axis 174, it has five associated GO BP terms: GO:0055072 (iron ion homeostasis), GO:0006879 (intracellular iron ion homeostasis), GO:0006121 (mitochondrial electron transport, succinate to ubiquinone), GO:0034553 (mitochondrial respiratory chain complex II assembly), and GO:0034552 (respiratory chain complex II assembly). The description of these terms suggests that the axis hallmarks the importance of iron ion homeostasis in mitochondrial respiration. Indeed, GO:0006121, GO:0034553, and GO:0034552 (average semantic similarity of 0.37) are connected with the mitochondrial respiratory chain. On the other hand, GO:0055072 and GO:0006879 (semantic similarity of 1) describe iron ion homeostasis To perform proper electron transfer in the mitochondrial respiratory chain, this organelle contains transition metals, and here iron is by far the most abundant [41]. Thus, proper homeostasis of iron ions is fundamental for cellular respiration.

Regarding axis 66, it has fifteen associated GO BP terms. These fifteen terms indicate that the axis is connected with the regulation of the cell's metabolism and growth via the TOR pathway activation. In particular, with an average semantic similarity of 0.91, GO:0006094 (gluconeogenesis), GO:0019319 (hexose biosynthetic process), and GO:0046364 (monosaccharide biosynthetic process) are related to sugar metabolism. On the other hand, with an average semantic similarity of 0.97, GO:0018209 (peptidyl-serine modification) and GO:0018105 (peptidyl-serine phosphorylation) are connected to the peptidyl-serine phosphorylation of proteins, which has been reported to be connected with the budding yeast central metabolism [42]. GO:0006808 describes also another process related to metabolism, such as the regulation of nitrogen utilization. Thus, up to now, all these GO BP terms are connected with functions related to the cell's metabolism. On the other hand, with a semantic similarity of 0.92, GO:0040008 (regulation of growth) and GO:0001558 (regulation of cell growth) are all related to cell growth. With an average semantic similarity of 0.57, GO:0030029 (actin filament-based process), GO:0030036 (actin cytoskeleton organization), GO:0030950 (establishment, or maintenance of actin cytoskeleton polarity), and GO:0030952 (establishment, or maintenance of cytoskeleton polarity) are connected to the actin cytoskeleton organization, which in turn is linked with the cell growth [43]. Regarding, GO:1905356 describes the regulation of snRNA pseudouridine synthesis, which is known to be modulated by the TOR signaling pathway being part of the growth program in budding yeast [44, 45]. Finally, GO:0031929 (TOR signaling) and GO:0031930 (mitochondria-nucleus signaling pathway) are connected with different signaling pathways. These pathways connect all the GO BP terms associated with the axes since it is known to affect the cell and its metabolism [46, 47].

With respect to axes that capture functional interactions that are not described in the literature, but are biologically coherent, we find axes 4 and 158.

Axis 158 has five associated GO BP terms. These five terms suggest that the axis is related to the cellular responses to changes in copper ion concentration. In particular, with a semantical similarity of 0.99, GO:0055070 (copper ion homeostasis) and GO:0006878 (intracellular copper ion homeostasis) describe copper ion homeostasis. The homeostasis of the copper ion involves several responses including epigenetic regulation, activation of DNA repair, or the production of proteins [48, 49]. Among these three responses, our results point out for the first time the role of the histone H3K79 methylation (GO:0034729), the global genome nucleotide-excision repair (GO:0070911), and the nucleolar large rRNA transcription (GO:0042790) in the homeostasis of the copper ion.

To conclude, axis 4 has ten associated GO BP terms. By analyzing these ten terms, we find that this axis describes different aspects of cell metabolism revealing a new connection between the mitochondrial group I introns and the regulation of energy production by alternative metabolic pathways in budding yeast. In particular, GO:0046942 and GO:0015718 (semantic similarity of 0.88) describe alternative metabolic pathways that use carbon sources such as carboxylic acids [50]. With a semantical similarity of 0.60, GO:0042407 (cristae formation) and GO:0045041 (protein import into mitochondrial intermembrane space), describe the architecture of the mitochondria. With an average pairwise semantic similarity of 0.66, GO:00196704 (NAD metabolic process), GO:0006116 (NADH oxidation), and GO:0006734 (NADH metabolic process) are linked to the REduction/OXidation (REDOX) reactions needed for the production of energy at the mitochondria. On the other hand, with an average semantic similarity of 0.55, GO:0090615 (mitochondrial mRNA processing), GO:0006316 (movement of group I intron), and GO:0006314 (intron homing), are connected with the mitochondrial group I introns. Mitochondrial introns are mobile genetic elements that form self-splicing RNA molecules [51]. These elements are divided into Group I and Group II introns depending on their secondary structure and splicing mechanism [52]. Group I introns encode other protein-coding genes in one of their loop regions including mitochondrial genes involved in the oxidative phosphorylation pathway [51]. Since the oxidative phosphorylation pathway has been related to the production of energy [51], we hypothesize that the group I introns may be key for the regulation of the cell's metabolism.

In conclusion, we demonstrate that our axes-based methodology captures new interactions between GO BP terms that are not described in the Gene Ontology. Moreover, we show that these interactions do not represent the functional similarity between GO BP terms as represented in the original Gene Ontology, but their functional interaction in higher-order cellular processes.

#### The Axes of the embedding space synthesize the core functions of different species' cells

In Results, section The axes of the embedding space capture the functional mechanisms of the cell of the main manuscript, we analyze the biological meaning of the ASFAs obtained from the axes of the human ONMTF embedding space. In particular, we show that ASFAs correctly summarize the biological information captured by the axes and describe coherent human cellular functions. Here, we further validate these observations by discussing more examples of human ASFAs. Then, we extend this analysis to the other five species: *Saccharomyces cerevisiae* (budding yeast), *Schizosaccharomyces pombe* (fission yeast), *Rattus norvegicus* (rat), *Drosophila melanogaster* (fruit fly) and *Mus musculus* (mouse).

Examples of human ASFAs include axis 12, which captures seven GO BP terms related to the regulation of cell adhesion (GO:0060354), leukocyte adhesion (GO:1904995), stem cell proliferation (GO:2000647), hematopoietic stem cell proliferation (GO:190233 and GO:1902034), and TRAIL-dependant apoptotic pathways (GO:1903121 and GO:1903122). The resulting ASFA combines and summarizes the keywords of these terms (endothelial, negative, regulation, apoptotic, molecule, signaling, cell, stem, activated, leukocyte, vascular, TRAIL, proliferation, adhesion, hematopoietic and production), representing a coherent function associated with the induction of apoptosis of tumor and infected cells via TNF-related apoptosis-inducing ligand (TRAIL) [53] (see Table 7). TRAIL also coordinates the immune response to tumor and infected cells by activating the leukocyte production by hematopoiesis and regulating inflammation [54].

Another example is the ASFA of axis 495 in human captures five GO BP terms that describe the response to endoplasmic unfolded protein (GO:1900101, GO:1903891 and GO:1903893) and the regulation of gene expression in response to cellular stress (GO:1990440 and GO:0036003). As can be seen in Table 7, its corresponding ASFA (polymerase II, mediated, RNA, unfolded, response, regulation, protein, stress, reticulum, ATF6, promoter, positive, transcription, endoplasmic) correctly summarizes these terms and displays a coherent biological function related to the cellular response against the accumulation of misfolded proteins in the Endoplasmic Reticulum [55].

Also, we extend this analysis to the other five species. To this aim, we generate the corresponding embedding spaces by applying ONMTF on the corresponding species PPI network (detailed in Methods, sections Biological datasets and Embedding the PPI networks of the main manuscript). We generate these embedding spaces with increasing dimensionalities (from 50 to 1000 dimensions with a step of 50). To select the optimal dimensionality of these embedding spaces, we follow the same criteria we did for the human PPI network embedding space (detailed in Supplementary Results, section Exploring the impact of the network embedding space's dimensionality on the biological information captured by the axes of the main manuscript). This dimensionality corresponds to 200, 200, 300, 250, and 400 for budding yeast, fission yeast, fruit fly, rat and mouse embedding spaces, respectively. Then, we use the GO BP terms captured by each axis to generate the ASFAs of each species (detailed in Methods, section Generating the Axes-Specific Functional Annotations of the main manuscript), and we analyze their biological coherence by literature curation.

We find that all the species ASFAs describe coherent functions of their corresponding species. For instance, the ASFA of axis 79 in budding yeast represents the trafficking of endosomes (see Supplementary Table 7). Curiously, this yeast is one of the most used models to study this transport process [56]. Another example in budding yeast is the ASFA of axis 82, which is connected to regulating gene expression via mRNA degradation (see Supplementary Table 7). Precisely with a process that involves the capping of the 7-methylguanosine residue that occurs after the deadenylation of the 3' poly(A) tracts of eukaryotic mRNAs and that serves as a backup mechanism to trigger mRNA decay if initial deadenylation is compromised [57]. Moreover, the ASFA of axis 20 in fission yeast is connected to the generation of large the ribosomal subunit necessary to synthesize proteins [58] (see Supplementary Table 7). Another example in this yeast is the ASFA of axis 32, which is also related to the synthesis of proteins (see Supplementary Table 7). However, in this case, the ASFA describes the regulation of protein synthesis via the rapamycin kinase complex I (TORC1) and II (TORC2). In the presence of ample nutrients, TORC1 and TORC2 activate and drive protein, lipid, and nucleotide synthesis by phosphorylating a wide range of proteins [59].

Regarding the fruit fly, we find ASFAs that represent functions that are more complex than the ones observed for the previous yeasts, such as the development of specific tissues. For instance, the ASFA of axis 1 describes the development of the visual nervous system (see Supplementary Table 7). Briefly, this tissue appears after the differentiation of the neuroectoderm by activating different epidermal growth factor receptors, such as ERBB2 [60]. Another example in the fruit fly is the ASFAs of axis 28, which is related to the wing imaginal disc of this fly (see Supplementary Table 7). This disc is a tissue of undifferentiated cells that are precursors of the wing and serves as a commonly used model system to study the regulation of growth [61].

Finally, we find that the ASFAs of mouse and rat are also connected to complex cellular functions, such as the immune system, or the nervous system. For instance, the ASFA of axis 41 of mouse describes the production of interferon-alpha, interleukins, and cytokines, during the cellular response to a virus infection [62] (see Supplementary Table 7). On the other hand, the ASFAs of axes 69 and 84 in rat, are connected to the synapsis of neurons and the production of steroids, respectively (see Supplementary Table 7).

In conclusion, these results demonstrate that the ASFAs describe coherent biological functions. The complete Tables with all the sets of species ASFAs can be found in the Supplementary online data.

#### The Axes of the embedding space give insights into the evolutionary history of species

In Results, section The axes of the embedding space uncover the human evolutionary history of the main manuscript, we show that the human ASFAs give insights into the evolutionary history of humans. In particular, we show that "prokaryotes" ASFAs reveal connections between complex human cellular functions to ancient prokaryote ones, "eukaryotes" ASFAs reveal evolutionary connections between humans and other eukaryotes and "vertebrate" ASFAs describe specific human traits that are unique to vertebrates. Here, we further validate these observations by discussing more examples of human ASFAs. Then, we extend this analysis to the ASFAs of the other five species: *Saccharomyces cerevisiae* (budding yeast), *Schizosaccharomyces pombe* (fission yeast), *Rattus norvegicus* (rat), *Drosophila melanogaster* (fruit fly) and *Mus musculus* (mouse).

Among the "prokaryotes" ASFAs in human, another example of ASFA that reveals connections between complex human cellular functions to ancient prokaryote ones is axis 61's ASFA (see Supplementary Table 7). This ASFA is extremely conserved (conservation degree of 20) and is connected with the RNA preprocessing by the spliceosome. Interestingly, although the origins of the spliceosome are debated, it is widely believed to have evolved from the ancestor of group II introns that emerged within bacteria during eukaryogenesis [63, 64].

On the other hand, "eukaryotes" ASFAs in human reveal evolutionary connections between humans and other eukaryotes. For instance, axis 120's ASFA describes a function related to the visual sense (see Supplementary Table 7). Among the taxons that are connected to this ASFA, we find mammals, such as mice (taxon id: 10090) and rats (taxon id: 10116), but also insects, such as the fruit fly (taxon id: 7227). Despite the divergence in the light receptors between these species, this axis further confirms that these receptors evolved from a common photoreceptor eukaryotic ancestor [65]. Also, the highest conservation degree among "eukaryotes" ASFAs is observed in axis 79. It describes the molecular mechanisms involved in the development of the heart and thyroid gland (see Supplementary Table 7). Species connected to this ASFA include animals that possess these organs, such as rats (taxon id: 10116), chickens (taxon id: 9031), and mice (taxon id: 10090), but also eukaryotes that lack these structures, including the budding yeast (taxon id: 559292), fission yeast (taxon id: 4896), and rice (taxon id: 39947). This suggests that the molecular mechanisms underlying the development of these organs originated early in eukaryotic evolution. Indeed, it is hypothesized that molecular pathways involved in human organogenesis appeared early in the evolution of multicellular organisms through the redeployment of components found in unicellular organisms [66, 67].

Finally, we explore the remaining "vertebrate" ASFAs that we do not discuss in the main manuscript and confirm that all of them describe specific traits that are unique to vertebrates. For instance, we find ten ASFAs that describe cellular functions related to the adaptive immune system, which is a system restricted to vertebrates [68, 69], for instance, lymphocyte proliferation and the activation of natural killer lymphocytes (see axes 473 and 370 in Table 7). Furthermore, we see ASFAs connected to different regulatory processes of the cell and to metabolic processes (see axes 58, 28, 452, 99, 91, 257, 166, 75, and 91 in Supplementary online data).

We extend this analysis to the other five species. To this end, we generate the corresponding embedding spaces by applying ONMTF on the corresponding species PPI network (detailed in Methods, sections Biological datasets and Embedding the PPI networks of the main manuscript). We generate these embedding spaces with increasing dimensionalities (from 50 to 1000 dimensions with a step of 50). To select the optimal dimensionality of these embedding spaces, we follow the same criteria we did for the human PPI network embedding space (detailed in Supplementary Results, section Exploring the impact of the network embedding space's dimensionality on the biological information captured by the axes of the main manuscript). This dimensionality corresponds to 200, 200, 300, 250, and 400 for budding yeast, fission yeast, fruit fly, rat and mouse embedding spaces, respectively. Then, we use the GO BP terms captured by each axis to generate the ASFAs of each species (detailed in Methods, section Generating the Axes-Specific Functional Annotations of the main manuscript). To investigate the link between these ASFAs and evolution, we divide the ASFAs of each species into three classes according to their conservation degree: "prokaryotes," "eukaryotes," and "vertebrates" (detailed in Methods, section Quantifying the evolution conservation degrees of the ASFA of the main manuscript). Then, We analyze in detail the meaning of these groups of ASFAs in the context of evolution.

We find that 78%, 69%, 59%, 63%, and 40% of all ASFAs in budding yeast, fission yeast, fruit fly, rat, and mouse, respectively are classified as "prokaryotes." These ASFAs present the lowest conservation degree in all the studied species, i.e., they are conserved in evolution (see Supplementary Figure 7). We observe that they represent the most basic molecular mechanisms of the cell, such as the translational process in budding yeast, the homeostasis of proteins in fission yeast, the homeostasis of ions in the fruit fly, or the lipid metabolism in mice (see axes 77, 57, 4, and 5, respectively in Supplementary Table 7).

On the other hand, we find that 22%, 31%, 41%, 33%, and 41% of all ASFAs in budding yeast, fission yeast, fruit fly, rat, and mouse, respectively are classified as "eukaryotes." These ASFAs have on average a lower conservation degree than the "prokaryotes" ones, i.e., they are newer in the evolutionary history. We find that they describe cellular functions that are connected to basic eukaryotic functions, e.g., with Golgi apparatus in budding yeast, signalling transduction in fission yeast, or cytoskeleton (see axes 79, 32, and 51, respectively in Supplementary Table 7).

Finally, as expected, the only organism that has "vertebrates" ASFAs are rat and mouse. Precisely, we find that 11% and 10% of all ASFAs are classified as "vertebrates" in rats and mice, respectively. These ASFAs have on average the lowest conservation degree, i.e., are the newest in evolution and they describe complex biological functions, such as estrous cycle, or odontogenesis in rats, and eyes' lens development, or blastocyst development in mice (see axes 81, 19, 86, and 80, respectively in Supplementary Table 7).

In conclusion, these results demonstrate that the ASFAs of different species can be used to give insights into their evolutionary history.

### Supplementary Figures

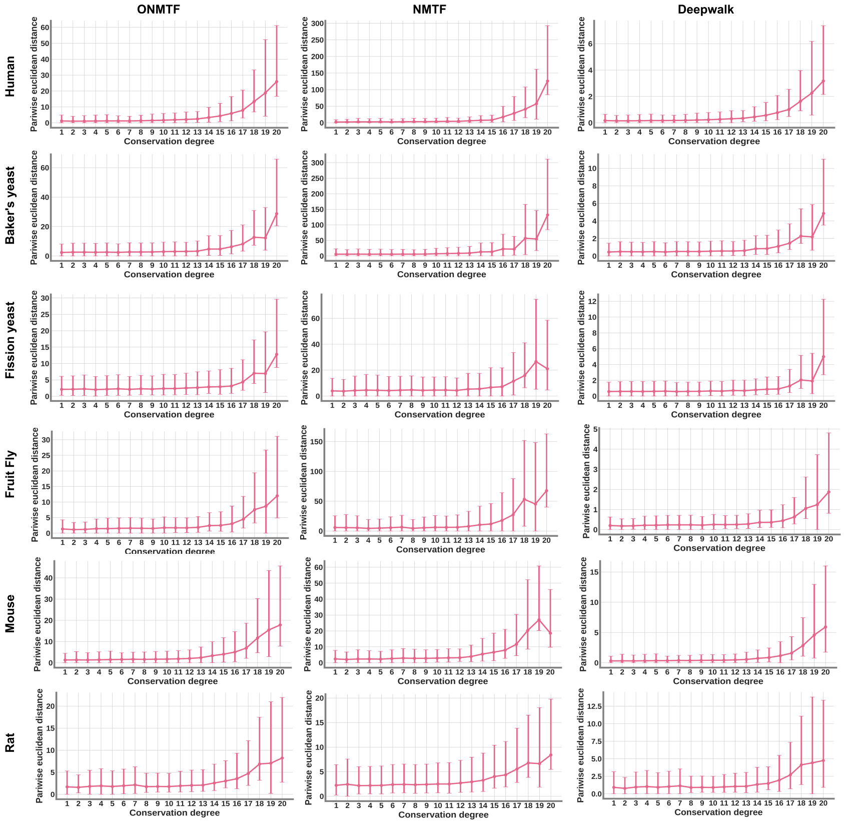


**Supplementary Figure 1**: The conservation degree of the GO BP terms influences the positions of their embedding vector in the species PPI network embedding space. We embed GO BP terms into the embedding spaces generated by applying ONMTF, NMTF, and Deepwalk algorithms on the species PPI network of *Homo sapiens sapiens* (denoted by human), *Saccharomyces cerevisiae* (denoted by budding yeast), *Schizosaccharomyces pombe* (denoted by fission yeast), *Rattus norvegicus* (denoted by rat), *Drosophila melanogaster* (denoted by fruit fly), and *Mus musculus* (denoted by mouse) (detailed in Methods, sections Biological datasets and Embedding the PPI networks of the main manuscript). We study the correlation between the mutual positions of their embedding vectors in the space (measured by their pairwise Euclidean distances) and their conservation degree (detailed in Methods, section Quantifying the evolutionary conservation of biological functions of the main manuscript). In each panel, the horizontal axis displays the conservation degree of the GO BP terms and the vertical axis shows the pairwise Euclidean distance distribution of their embedding vectors.

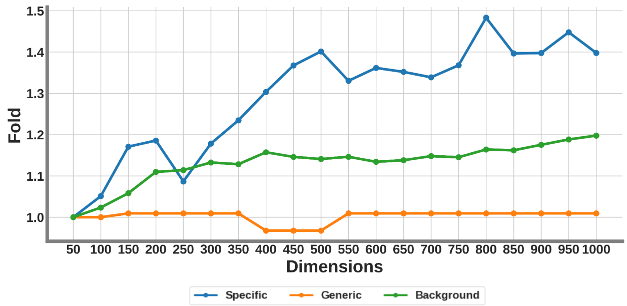


**Supplementary Figure 2**: Specific biological functions are captured by the axes of the human ONMTF embedding spaces with the increment of dimensions. We take as reference the lowest dimensional embedding space (50 dimensions) and compare the fold increase between the number of “specific,” “generic,” and “background” GO BP terms associated with its axes and with those captured by the axes of the subsequent species PPI network embedding spaces. The horizontal axis displays the number of dimensions of the embedding space.

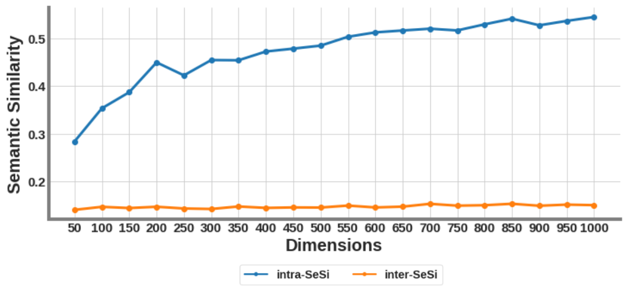


**Supplementary Figure 3**: Increasing the dimensionality enhances the stratification of the biological information captured by the axes of the human ONMTF embedding spaces. We compute Lin’s semantic pairwise semantic similarity between any two GO BP terms. The blue line shows the average semantic similarity of the pairs of GO BP terms that are associated with the same axis (intra-SeSi). The orange line shows the average semantic similarity of the pairs of GO BP terms that are associated with different axis (inter-SeSi). The horizontal axis displays the number of dimensions of the embedding space.

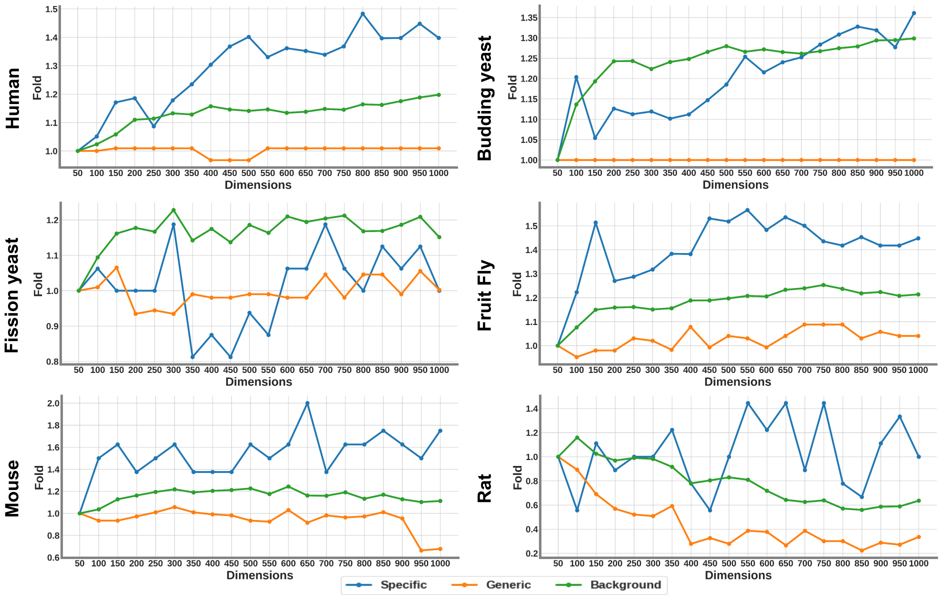


**Supplementary Figure 4**: Specific biological functions are captured by the axes of the species ONMTF embedding spaces with the increment of dimensions. We generate the species PPI network embedding spaces by applying ONMTF on the species PPI network of *Homo sapiens sapiens* (denoted by human), *Saccharomyces cerevisiae* (denoted by budding yeast), *Schizosaccharomyces pombe* (denoted by fission yeast), *Rattus norvegicus* (denoted by rat), *Drosophila melanogaster* (denoted by fruit fly), and *Mus musculus* (denoted by mouse) (detailed in Methods, sections Biological datasets and Embedding the PPI networks of the main manuscript). We generate these embedding spaces with increasing dimensionalities (from 50 to 1000 dimensions with a step of 50). For each species embedding space, we take as a reference the 50-dimensional embedding space and we compute the fold between the number of "specific," "generic," and "background" functional annotations associated with its axes and that of the subsequent species PPI network embedding spaces (detailed in Methods, section Quantifying the evolutionary conservation of biological functions of the main manuscript, and Results, section Exploring the impact of the network embedding space's dimensionality on the biological information captured by the axes of the main manuscript). The horizontal axis displays the number of dimensions of the embedding space. The horizontal axis displays the number of dimensions of the embedding space.

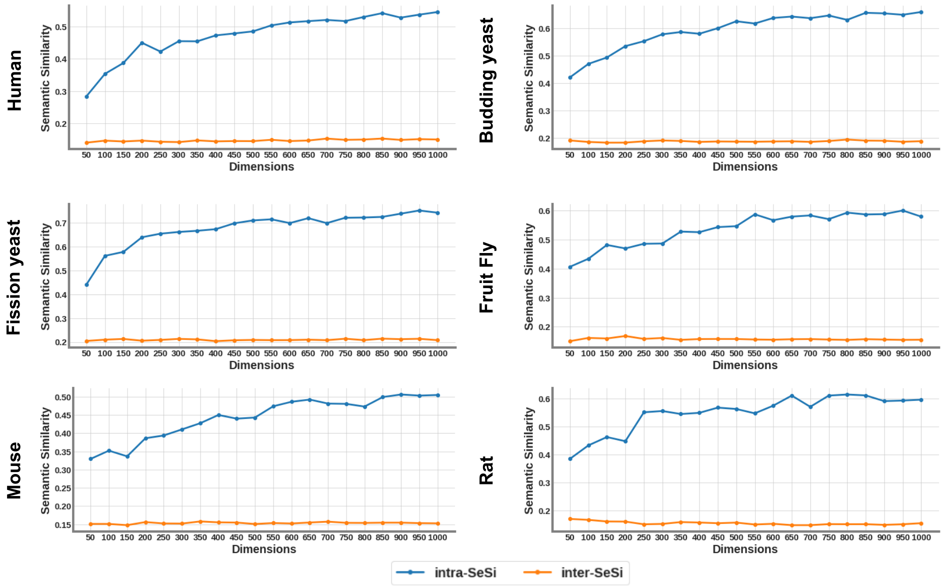


**Supplementary Figure 5**: Specific biological functions are disentangled by the axes of the species ONMTF embedding spaces with the increment of dimensions. We generate the species PPI network embedding spaces by applying ONMTF on the species PPI network of *Homo sapiens sapiens* (denoted by human), *Saccharomyces cerevisiae* (denoted by budding yeast), *Schizosaccharomyces pombe* (denoted by fission yeast), *Rattus norvegicus* (denoted by rat), *Drosophila melanogaster* (denoted by fruit fly), and *Mus musculus* (denoted by mouse) (detailed in Methods, sections Biological datasets and Embedding the PPI networks of the main manuscript). We generate these embedding spaces with increasing dimensionalities (from 50 to 1000 dimensions with a step of 50). For each species embedding space, we compute Lin's semantic pairwise semantic similarity between any two GO BP terms (detailed in Methods, section Quantifying the evolutionary conservation of biological functions of the main manuscript). The blue line shows the average semantic similarity of the pairs of GO BP terms that are associated with the same axis (intra-SeSi). The orange line shows the average semantic similarity of the pairs of GO BP terms that are associated with different axis (inter-SeSi). The horizontal axis displays the number of dimensions of the embedding space.

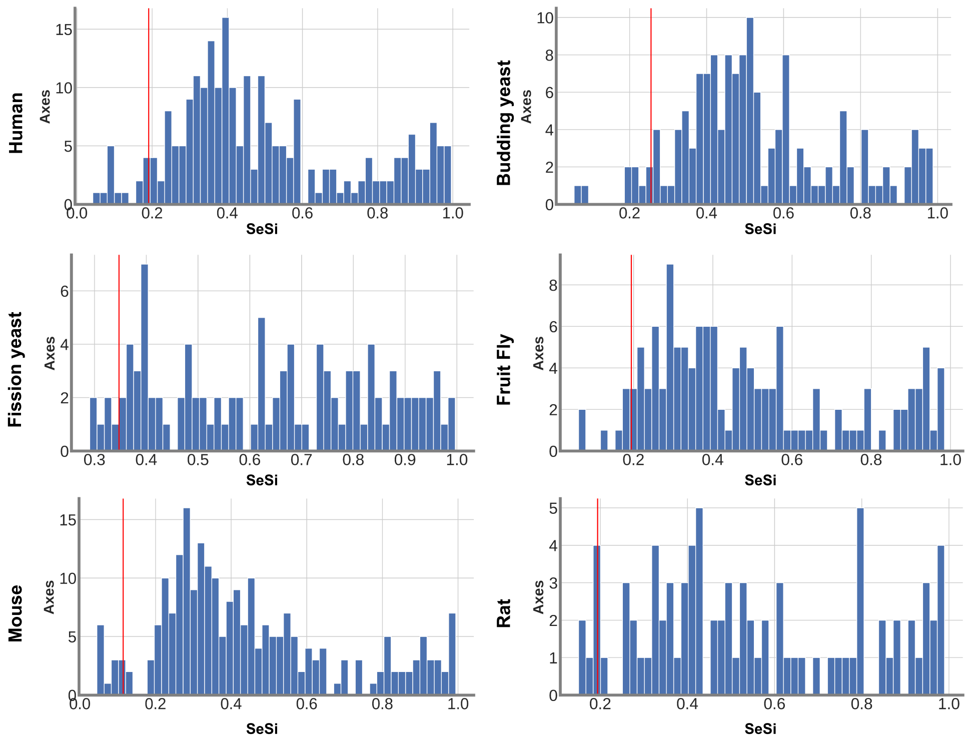


**Supplementary Figure 6**: Distribution of "intra-axis SeSi." We generate the species PPI network embedding spaces by applying ONMTF on the species PPI network of *Homo sapiens sapiens* (denoted by human), *Saccharomyces cerevisiae* (denoted by budding yeast), *Schizosaccharomyces pombe* (denoted by fission yeast), *Rattus norvegicus* (denoted by rat), *Drosophila melanogaster* (denoted by fruit fly), and *Mus musculus* (denoted by mouse) (detailed in Methods, sections Biological datasets and Embedding the PPI networks of the main manuscript). We generate these embedding spaces with 500, 200, 200, 250, 300, and 400 dimensions, respectively, since these dimensionalities correspond to the optimal dimensionality of such spaces (as detailed in section 3 of the main text). For each species embedding space, we compute Lin's semantic pairwise semantic similarity between any two GO BP terms (detailed in Methods, section Quantifying the evolutionary conservation of biological functions of the main manuscript). Then, for each axis, we report the average Lin's semantic pairwise semantic similarity of the pairs of GO BP terms that are associated with it (that we term "intra-axis SeSi"). For each panel, the horizontal axis ("SeSi") show the "intra-axis SeSis" and the vertical axis ("Axes") the number of embedding axes with a certain "intra-axis semantic similarities." The red line represents the $5^{th}$ percentile of the distributions. We use this threshold to define the set of axes that captures the largest number of new functional interactions between the GO BP terms (detailed in Results, section The axes of the embedding space uncover new functional interactions between GO BP terms of the main text).

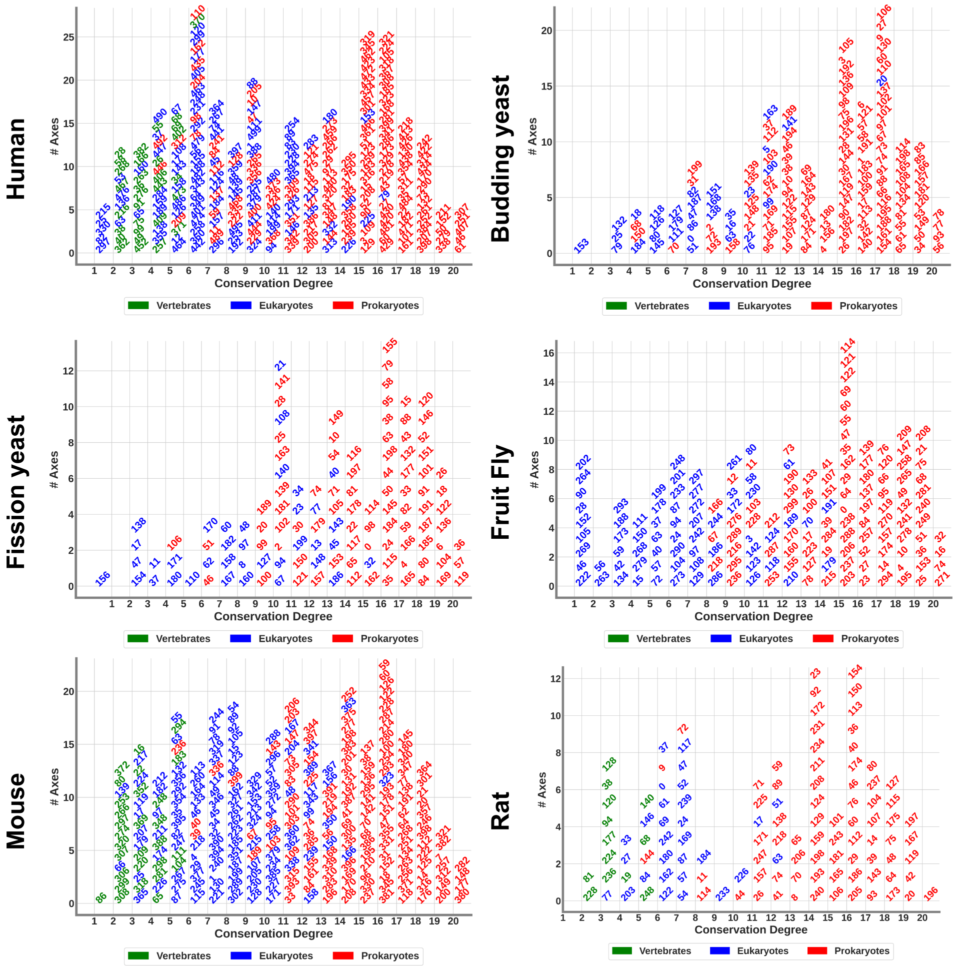


**Supplementary Figure 7**: The ASFAs give insights into the evolutionary history of *Saccharomyces cerevisiae* (denoted by budding yeast), *Schizosaccharomyces pombe* (denoted by fission yeast), *Rattus norvegicus* (denoted by rat), *Drosophila melanogaster* (denoted by fruit fly), and *Mus musculus* (denoted by mouse). For each species, we use the conservation degree of its ASFAs to divide them into three groups: "prokaryotes," "eukaryotes," and "vertebrates" (detailed in Methods, section Quantifying the evolution conservation degrees of the ASFA of the main manuscript). Then, we order the ASFAs according to their conservation degree. In each panel, the horizontal axis displays the conservation degree of the ASFAs and the vertical axis shows the number of ASFAs with a certain conservation degree. Each ASFA is represented in the panels by the number of the axis from which it was obtained.

### Supplementary Tables

| Network | #Nodes | #Edges | #Density |
|---|---|---|---|
| Human | 18,290 | 368,180 | 0.0022 |
| Budding yeast | 5,887 | 111,307 | 0.0064 |
| Fission yeast | 3,269 | 10,958 | 0.0020 |
| Fruit fly | 8,917 | 49,756 | 0.0012 |
| Mouse | 8,043 | 26,661 | 0.0008 |
| Rat | 2,847 | 5,252 | 0.0013 |

**Supplementary Table 1**: The statistics of the species PPI networks. For the six species: *Homo sapiens sapiens* (denoted by "Human"), *Saccharomyces cerevisiae* (denoted by "Budding yeast"), *Schizosaccharomyces pombe* (denoted by "Fission yeast"), *Drosophila melanogaster* (denoted by "Fruit fly"), *Mus musculus* (denoted by "Mouse") and *Rattus norvegicus* (denoted by "Rat"). The first column, "Network," lists the species. The second column "# Nodes," show the number of nodes in the species PPI network. The third column, "# Edges," contains the number of edges between the nodes. The fourth column, "# Density," specifies the edge density of the corresponding species PPI network.

| Species | # GO BP terms |
|---|---|
| Human | 6,864 |
| Budding yeast | 3,042 |
| Fission yeast | 1,864 |
| Fruit fly | 3,712 |
| Rat | 2,828 |
| Mouse | 6,343 |

**Supplementary Table 2**: Number of GO BP annotations for each species PPI network. For the six species: *Homo sapiens sapiens* (denoted by "Human"), *Saccharomyces cerevisiae* (denoted by "Budding yeast"), *Schizosaccharomyces pombe* (denoted by "Fission yeast"), *Drosophila melanogaster* (denoted by "Fruit fly"), *Rattus norvegicus* (denoted by "Rat") and *Mus musculus* (denoted by "Mouse"). The first column, "Network," lists the species. The second column, "# GO BP terms," presents the number of GO BP terms that annotates at least one gene in the corresponding species PPI network.

| Embedding algorithm | % Axes | % GO |
|---|---|---|
| ONMTF | 53.72 | 57.40 |
| NMTF | 61.80 | 48.12 |
| Deepwalk | 68.00 | 35.50 |

**Supplementary Table 3**: On average, the axes of the species PPI network embedding spaces generated by the ONMTF embedding algorithm are the best for capturing the cell's functional organization from PPI networks. We generate the species PPI network embedding spaces by applying ONMTF, NMTF, and Deepwalk algorithms on the species PPI network of *Homo sapiens sapiens*, *Saccharomyces cerevisiae*, *Schizosaccharomyces pombe*, *Rattus norvegicus*, *Drosophila melanogaster*, and *Mus musculus* (detailed in Methods, sections Biological datasets and Embedding the PPI networks). We generate these embedding spaces with increasing dimensionalities (from 50 to 1000 dimensions with a step of 50). For each species PPI network embedding space, we use our new axes-based method to capture the GO BP terms that we embed in the space (detailed in Methods, section Annotating the axes of the gene embedding space with GO BP terms of the main manuscript). The first column, "Embedding algorithm," lists the embedding algorithms used for generating the embedding spaces. The second column, " % Axes," presents the percentage of axes that captures at least one embedded GO BP term averaged across dimensions and species. The third column, " % GO," shows the percentage of the total embedded GO BP terms that are associated with the axes of the space averaged across dimensions and species.

| Embedding algorithm | Intra SeSi | Inter SeSi | Random SeSi | Shortest Paths |
|---|---|---|---|---|
| ONMTF | 0.50 | 0.16 | 0.16 | 3.71 |
| NMTF | 0.42 | 0.16 | 0.16 | 3.90 |
| Deepwalk | 0.35 | 0.16 | 0.16 | 4.31 |

**Supplementary Table 4**: On average, the GO BP terms captured by the axes of the human PPI network embedding spaces generated by the ONMTF embedding algorithm are more coherent and better organized than those of the NMTF and Deepwalk spaces. We generate the human PPI network embedding spaces by applying ONMTF, NMTF, and Deepwalk algorithms on the PPI network of *Homo sapiens sapiens* (detailed in Methods, sections Biological datasets and Embedding the PPI networks). We generate these embedding spaces with increasing dimensionalities (from 50 to 1000 dimensions with a step of 50). For each human PPI network embedding space, we use our new axes-based method to capture the GO BP terms that we embed in the space (detailed in Methods, section Annotating the axes of the gene embedding space with GO BP terms). Then, we investigate how coherently the captured GO BP terms are distributed across the axes according to the Gene Ontology (detailed in Methods, section Assessing the ability of our axes-based methodology to capture biological information of the main manuscript). The first column, "Embedding algorithm," lists the embedding algorithms used for generating the embedding spaces. The second column, "Intra SeSi," shows the average Lin's semantic similarity between the GO BP terms that are associated by the same axis averaged across dimensions and species. The third column, "Inter SeSi," presents the average Lin's semantic similarity between the GO BP terms that are captured by different axes averaged across dimensions and species. The fourth column, "Random SeSi," shows the global average Lin's semantic similarity between any two GO BP terms. The fifth column, "Shortest Paths," displays the mean shortest paths in the GO ontology-directed acyclic graph between the GO BP terms associated with the same axis averaged across dimensions and species.

| Embedding algorithm | Intra SeSi | Inter SeSi | Random SeSi | Shortest Paths |
|---|---|---|---|---|
| ONMTF | 0.54 | 0.16 | 0.16 | 3.71 |
| NMTF | 0.48 | 0.18 | 0.16 | 3.90 |
| Deepwalk | 0.46 | 0.18 | 0.16 | 4.31 |

**Supplementary Table 5**: On average, the GO BP terms captured by the axes of the species PPI network embedding spaces generated by the ONMTF embedding algorithm are more coherent and better organized than those of the NMTF and Deepwalk spaces. We generate the species PPI network embedding spaces by applying ONMTF, NMTF, and Deepwalk algorithms on the species PPI network of *Homo sapiens sapiens*, *Saccharomyces cerevisiae*, *Schizosaccharomyces pombe*, *Rattus norvegicus*, *Drosophila melanogaster*, and *Mus musculus* (detailed in sections Biological datasets and Embedding the PPI networks of the main manuscript). We generate these embedding spaces with increasing dimensionalities (from 50 to 1000 dimensions with a step of 50). For each species PPI network embedding space, we use our new axes-based method to capture the GO BP terms that we embed in the space (detailed in Methods, section Annotating the axes of the gene embedding space with GO BP terms). Then, we investigate how coherently the captured GO BP terms are distributed across the axes according to the Gene Ontology (detailed in Methods, section Assessing the ability of our axes-based methodology to capture biological information of the main manuscript). The first column, "Embedding algorithm," lists the embedding algorithms used for generating the embedding spaces. The second column, "Intra SeSi," shows the average Lin's semantic similarity between the GO BP terms that are associated by the same axis averaged across dimensions and species. The third column, "Inter SeSi," presents the average Lin's semantic similarity between the GO BP terms that are captured by different axes averaged across dimensions and species. The fourth column, "Random SeSi," shows the global average Lin's semantic similarity between any two GO BP terms. The fifth column, "Shortest Paths," displays the mean shortest paths in the GO ontology-directed acyclic graph between the GO BP terms associated with the same axis averaged across dimensions and species.

| Species | # Dimensions |
|---|---|
| Human | 500 |
| Budding yeast | 200 |
| Fission yeast | 200 |
| Fruit fly | 300 |
| Rat | 250 |
| Mouse | 400 |

**Supplementary Table 6**: The optimal number of dimensions for the six species ONMTF embedding spaces. For the species PPI network embedding spaces generated by applying the ONMTF algorithm on the species PPI network of *Homo sapiens sapiens* (denoted by human), *Saccharomyces cerevisiae* (denoted by budding yeast), *Schizosaccharomyces pombe* (denoted by fission yeast), *Rattus norvegicus* (denoted by rat), *Drosophila melanogaster* (denoted by fruit fly), and *Mus musculus* (denoted by mouse), we use our axes-based method to find their optimal dimensionality (detailed in Results, section Exploring the impact of the network embedding space's dimensionality on the biological information captured by the axes of the main manuscript). The first column, "Species," lists the species. The second column, "# Dimensions," shows the optimal dimensionality of the species PPI network embedding space according to our axes-based method.

| Species | Axis | Terms | #GO | Taxons |
|---|---|---|---|---|
| Human | 12 | endothelial, negative, regulation, apoptotic, molecule, signaling, cell, stem, activated, leukocyte, vascular, TRAIL, proliferation, adhesion, hematopoietic, production | 7 | 7227, 7955, 9606, 10090, 10116 |
| Human | 61 | subunit, spliceosome, processing, nucleobase, RNA, aromatic, heterocycle, snRNP, complex, compound, process, capping, nucleophile, assembly, containing, reactions, spliceosomal, cellular, 3', mRNA, adenosine, ribonucleoprotein, organization, organic, cyclic, bulged, transesterification, splicing, nucleic, metabolic | 20 | 3702, 4896, 6239, 7227, 7955, 9031, 9606, 9615, 9823, 9913, 10090, 10116, 36329, 39947, 195103, 214684, 227321, 352472, 511145, 559292 |
| Human | 79 | heart, thyroid gland, organ, anatomical development | 5 | 10116, 9031, 10090, 9823, 7955, 7227, 6239, 9606, 4896, 214684, 352472, 559292, 227321, 39947, 3702, 352472 |
| Human | 473 | negative, regulation, activation, cell, proliferation, lymphocyte | 3 | 9031, 9606, 9913, 10090, 10116 |
| Human | 370 | mediated, natural, killer, leukocyte, activation, cytotoxicity, immunity, lymphocyte, cell, activation | 6 | 7955, 9606, 9615, 9823, 10090, 10116 |
| Human | 120 | system, light, visual, nervous, stimulus, process, sensory, perception | 4 | 6239, 7227, 7955, 9606, 10090, 10116 |
| Budding yeast | 79 | endosome, Golgi, early, transport, | 1 | 559292, 9606, 6239 |
| Budding yeast | 82 | decapping, methylguanosine, RNA, cap, nuclear, deadenylation, mRNA, dependent, transcribed | 3 | 4896, 9606, 10090, 3702, 7227, 559292, 6239 |
| Budding yeast | 77 | methylation, subunit, benzene, regulation, translation, nucleus, initiation, compound, fidelity, process, gene, export, rRNA, assembly, amide, expression, small, containing, tRNA, positive, transport, post-transcriptional, ribosomal, cellular, translational, metabolic | 12 | 4896, 214684, 10116, 9606, 9031, 511145, 10090, 36329, 39947, 195103, 9615, 7955, 3702, 352472, 9913, 7227, 559292, 227321, 6239, 9823 |
| Fission yeast | 20 | subunit, large, biogenesis, complex, ribosomal, ribonucleoprotein | 2 | 4896, 9606, 36329, 10090, 511145, 7955, 3702, 7227, 559292 |
| Fission yeast | 32 | TORC2, regulation, TORC1, reproductive, signaling, process, positive | 6 | 4896, 10116, 9606, 9031, 10090, 9615, 7955, 3702, 352472, 9913, 7227, 559292, 227321, 6239, 9823 |
| Fission yeast | 57 | catabolic, protein, removal, conjugation, organonitrogen, compound, process, deneddylation, small, cellular, SCF, proteasomal, dependent, proteolysis, ubiquitin, metabolic, modification | 9 | 4896, 214684, 10116, 9606, 9031, 36329, 10090, 511145, 195103, 39947, 9823, 9615, 7955, 3702, 352472, 9913, 7227, 559292, 227321, 6239 |
| Fruit fly | 1 | synaptic, olfactory, mediated, vesicle, follicular, factor, negative, tyrosine, peptidyl, regulation, photoreceptor, epithelium, clathrin, filament, neuron, dorsal-ventral, transduction, eye, specification, signaling, cell, compound, commitment, learning, epidermal, growth, ERBB2, assembly, pathway, positive, transport, cascade, communication, dependent, organization, phosphorylation, fate, signal, modification, receptor | 25 | 4896, 10116, 9606, 9031, 511145, 10090, 9823, 9615, 7955, 3702, 352472, 9913, 7227, 559292, 6239 |
| Fruit fly | 28 | negative, vein, regulation, disc, derived, specification, imaginal, wing | 1 | 7227 |
| Fruit fly | 4 | cation, biosynthetic, metal, divalent, regulation, retinal, aldehyde, ion, compound, lipid, process, olefinic, inorganic, transport, diterpenoid, cellular, retinoid, homeostasis, metabolic | 9 | 4896, 214684, 10116, 9606, 9031, 511145, 10090, 36329, 39947, 9823, 9615, 7955, 3702, 352472, 9913, 7227, 559292, 6239 |
| Mouse | 41 | immune, type, lipopolysaccharide, negative, alpha, interferon, response, regulation, innate, pattern, signaling, involved, recognition, pathway, virus, dsRNA, interleukin, inflammatory, cytokine, production, receptor | 10 | 10116, 9606, 9031, 511145, 10090, 9823, 9615, 7955, 3702, 352472, 9913, 7227, 559292, 6239 |
| Mouse | 7 | biosynthetic, estrogen, glycerophospholipid, glycerolipid, lipid, process, phosphatidylcholine, metabolic | 5 | 4896, 214684, 10116, 9606, 9031, 36329, 10090, 511145, 39947, 195103, 9823, 9615, 7955, 3702, 352472, 9913, 7227, 559292, 6239 |
| Mouse | 86 | type, induction, lens, eye, camera | 1 | 10090 |
| Mouse | 80 | blastocyst, development | 1 | 10090, 9606 |
| Rat | 69 | synaptic, signaling, trans, anterograde, transmission, chemical | 4 | 10116, 9606, 10090, 7955, 7227, 6239 |
| Rat | 84 | mediated, intracellular, signaling, steroid, pathway, hormone, androgen, receptor | 3 | 10116, 9606, 10090, 3702, 7227 |
| Rat | 51 | negative, regulation, polymerization, ion, microtubule, polymerization, import, calcium | 4 | 4896, 10116, 9606, 9031, 10090, 7955, 3702, 352472, 9913, 7227, 559292, 6239 |
| Rat | 81 | estrous, cycle, ovulation | 2 | 10090, 10116 |
| Rat | 19 | odontogenesis | 1 | 10090, 7955, 10116, 9606 |

**Supplementary Table 7**: The species ASFAs describe coherent functions of six species. For the species PPI network embedding spaces generated by applying the ONMTF algorithm on the species PPI network of *Homo sapiens sapiens* (denoted by human), *Saccharomyces cerevisiae* (denoted by budding yeast), *Schizosaccharomyces pombe* (denoted by fission yeast), *Rattus norvegicus* (denoted by rat), *Drosophila melanogaster* (denoted by fruit fly), and *Mus musculus* (denoted by mouse), we use our new axes-based method to capture the GO BP terms that we embed in the space (detailed in Methods, sections Biological datasets, Embedding the PPI networks, and Annotating the axes of the gene embedding space with GO BP terms of the main manuscript). Then, we use the GO BP terms captured by the axes of the embedding spaces to generate the ASFAs (detailed in Methods, section Generating the Axes-Specific Functional Annotations of the main manuscript). The first column, "Species," lists the species. The second column, "Axis," lists the name of the axes from which each ASFA was obtained. The third column, "Terms," shows the description of the ASFAs. The fourth column, "#GO," displays the number of GO BP terms that are associated with the axis. The fifth column, "Taxons," shows the Taxonomy ID of the different species for which the associated GO BP terms appear. The complete Tables for all the species ASFAs can be found in the Supplementary online data.

| Species | Empty vs Non-Empty | Empty vs Random | Annotated Genes (empty axes) | Annotated Genes (non-empty axes) |
|---|---|---|---|---|
| Human | $1.76 \times 10^{-63}$ | $6.46 \times 10^{-28}$ | 40.0% | 53.8% |
| Budding yeast | $1.67 \times 10^{-8}$ | $6.94 \times 10^{-43}$ | 66.7% | 87.2% |
| Fission yeast | $7.42 \times 10^{-17}$ | $1.81 \times 10^{-62}$ | 23.7% | 61.4% |
| Fruit fly | $8.85 \times 10^{-15}$ | $1.05 \times 10^{-42}$ | 32.1% | 51.9% |
| Rat | 0.11 | $7.89 \times 10^{-6}$ | 51.7% | 76.1% |
| Mouse | 0.01 | $9.18 \times 10^{-30}$ | 52.5% | 74.5% |

**Supplementary Table 8**: Genes that are associated with the empty axes tend to form densely connected neighborhoods in the species PPI networks. We generate the species embedding spaces by applying the ONMTF algorithm on the species PPI network of *Homo sapiens sapiens* (denoted by human), *Saccharomyces cerevisiae* (denoted by budding yeast), *Schizosaccharomyces pombe* (denoted by fission yeast), *Rattus norvegicus* (denoted by rat), *Drosophila melanogaster* (denoted by fruit fly), and *Mus musculus* (denoted by mouse). For each species PPI network embedding space, we associate genes with their embedding axes. Then, we evaluate the connectivity in the original species PPI network by computing the clustering coefficient between genes associated with the same axis (detailed in Methods, section Generating the Axes-Specific Functional Annotations of the main manuscript). The first column, "Species," lists the species. The second column, "Empty vs Non-Empty," shows the p-value from a one-sided Mann-Whitney U test comparing if the clustering coefficient of the genes associated with non-empty axes (axes with at least one associated GO BP term) is statistically higher than the clustering coefficient of genes associated with empty axes (axes with non-associated GO BP terms). The third column, "Empty vs Random," displays the p-value from a one-sided Mann-Whitney U test comparing if the clustering coefficient of the genes associated with empty axes is statistically higher than expected by random. The fourth column, "Annotated Genes (empty axes)," displays the percentage of genes associated with empty axes that are annotated with at least one GO BP term. The fifth column, "Annotated Genes (non-empty axes)," shows the percentage of genes associated with non-empty axes that are annotated with at least one GO BP term.

| Axis | Terms | #Genes | Empty |
|---|---|---|---|
| 9 | neurotransmission, glycinergic, gonadotropin, unsaturated, choline, activating, glycosylation, glycine, adenylate, cyclase | 19 | Yes |
| 76 | chylomicron, brood, thymocyte, folding, transcription, microtubule, polymerase, leukocyte, helper, thymus | 27 | Yes |
| 68 | transcription, somitogenesis, polymerase, developmental, skeletal, commitment, midbrain, development, binding, dopaminergic | 78 | No |
| 370 | natural, killer, immunoglobulin, zinc, biosynthesis, transamidation, glutaminyl-tRNAGln, cytotoxicity, eye, adhesion | 20 | No |

**Supplementary Table 9**: The empty axes of the human ONMTF embedding space capture human cellular functions. For the human ONMTF embedding space, we use the genes associated with its empty axes (axes without associated GO BP terms) and non-empty axes to generate the ASFAs (detailed in Methods, section Generating the Axes-Specific Functional Annotations of the main manuscript). The first column, "Axis," lists the name of the axes from which each ASFA was obtained. The second column, "Terms," shows the description of the ASFAs (due to the length of these ASFAs, we show the top 10 words with the highest TF-IDF, i.e., the most relevant, see the complete ASFA in Supplementary online data). The third column, "#Genes," displays the number of genes that are associated with the axis. The fourth column, "Empty," indicates if the axis is empty ("Yes"), or not ("No"). The complete Table with all the human ASFAs generated by using the genes associated with its axes can be found in the Supplementary online data.

| Embedding algorithm | % Genes | % GO | % Clusters |
|---|---|---|---|
| ONMTF | 47.59 | 21.47 | 50.93 |
| NMTF | 35.98 | 20.58 | 49.18 |
| Deepwalk | 58.30 | 15.29 | 34.37 |

**Supplementary Table 10**: The species PPI network embedding spaces generated by the ONMTF are the best at capturing the cell's functional organization. We generate the species PPI network embedding spaces by applying ONMTF, NMTF, and Deepwalk algorithms on the species PPI network of *Homo sapiens sapiens*, *Saccharomyces cerevisiae*, *Schizosaccharomyces pombe*, *Rattus norvegicus*, *Drosophila melanogaster*, and *Mus musculus* (detailed in Methods, sections Biological datasets and Embedding the PPI networks of the main manuscript). We generate these embedding spaces with increasing dimensionalities (from 50 to 1000 dimensions with a step of 50). For each embedding space, we cluster genes whose embedding vectors are close in the space, and then we measure the enrichment of those clusters in GO BP annotations (detailed in the Supplementary Results, section Our axes-based method outperforms the classic gene-centric approach in capturing the cell's functional organization). The first column, "Embedding algorithm," lists the embedding algorithms used for generating the embedding spaces. The second column, "% Genes," presents the percentage of enriched genes in the clusters (out of the total number of genes in the corresponding species PPI network) averaged across dimensions and species. The third column, "% GO," shows the percentage of GO BP terms enriched in the clusters (out of the total number of GO BP terms) averaged across dimensions and species. The fourth column, "% Clusters," displays the percentage of clusters with at least one gene enriched (out of the total number of clusters) averaged across dimensions and species.

| Embedding algorithm | Fold | SeSi Axes | SeSi Clusters |
|---|---|---|---|
| ONMTF | 1.32 | 0.50 | 0.35 |
| NMTF | 1.23 | 0.42 | 0.38 |
| Deepwalk | 3.28 | 0.35 | 0.30 |

**Supplementary Table 11**: Our axes-based methodology outperforms the standard state-of-the-art gene-centric method in capturing biological from human PPI network embedding spaces. We generate the human PPI network embedding spaces by applying ONMTF, NMTF, and Deepwalk algorithms on the PPI network of *Homo sapiens sapiens* (detailed in Methods, sections Biological datasets and Embedding the PPI networks of the main manuscript). We generate these embedding spaces with increasing dimensionalities (from 50 to 1000 dimensions with a step of 50). For each embedding space, we apply our axes-based method and the standard gene-centric approach (detailed in Methods, section Assessing the ability of our axes-based methodology to capture biological information of the main manuscript and Supplementary Results, section Our axes-based method outperforms the classic gene-centric approach in capturing the cell's functional organization, respectively) and compare the GO BP terms captured by each method. The first column, "Embedding algorithm," lists the embedding algorithms used for generating the embedding spaces. The second column, "Fold," shows the fold between the different GO BP terms captured by our method and that for the gene-centric approach averaged across dimensions. The third column, "SS Axes," displays Lin's semantic similarity between GO BP terms associated with the same axis averaged across dimensions. The fourth column, "SS Clusters," shows Lin's semantic similarity between GO BP terms enriched in the same gene cluster averages across dimensions.

| Embedding algorithm | Fold | SeSi Axes | SeSi Clusters |
|---|---|---|---|
| ONMTF | 1.79 | 0.54 | 0.40 |
| NMTF | 1.10 | 0.48 | 0.40 |
| Deepwalk | 1.90 | 0.46 | 0.45 |

**Supplementary Table 12**: Our axes-based methodology outperforms the standard state-of-the-art gene-centric method in capturing biological from different species PPI network embedding spaces. We generate the species PPI network embedding spaces by applying ONMTF, NMTF, and Deepwalk algorithms on the species PPI network of *Homo sapiens sapiens*, *Saccharomyces cerevisiae*, *Schizosaccharomyces pombe*, *Rattus norvegicus*, *Drosophila melanogaster*, and *Mus musculus* (detailed in sections Biological datasets and Embedding the PPI networks of the main manuscript). We generate these embedding spaces with increasing dimensionalities (from 50 to 1000 dimensions with a step of 50). For each species embedding space, we apply our axes-based method and the standard gene-centric approach (detailed in Methods, section Assessing the ability of our axes-based methodology to capture biological information of the main manuscript and Supplementary Results, section Our axes-based method outperforms the classic gene-centric approach in capturing the cell's functional organization, respectively) and compare the GO BP terms captured by each method. The first column, "Embedding algorithm," lists the embedding algorithms used for generating the embedding spaces. The second column, "Fold," shows the fold between the different GO BP terms captured by our method and that for the gene-centric approach averaged across dimensions and species. The third column, "SS Axes," displays Lin's semantic similarity between GO BP terms associated with the same axis averaged across dimensions and species. The fourth column, "SS Clusters," shows Lin's semantic similarity between GO BP terms enriched in the same gene cluster averaged across dimensions and species.

| Embedding algorithm | Rand | SeSi Axes | SeSi Clusters |
|---|---|---|---|
| ONMTF | 0.14 | 0.45 | 0.31 |
| NMTF | 0.08 | 0.33 | 0.31 |
| Deepwalk | 0.04 | 0.45 | 0.35 |

**Supplementary Table 13**: Our axes-based methodology captures biological information from different species PPI network embedding spaces that differs from that captured by the state-of-the-art gene-centric method. We generate the species PPI network embedding spaces by applying ONMTF, NMTF, and Deepwalk algorithms on the species PPI network of *Homo sapiens sapiens*, *Saccharomyces cerevisiae*, *Schizosaccharomyces pombe*, *Rattus norvegicus*, *Drosophila melanogaster*, and *Mus musculus* (detailed in Methods, sections Biological datasets and Embedding the PPI networks of the main manuscript). We generate these embedding spaces with increasing dimensionalities (from 50 to 1000 dimensions with a step of 50). For each species embedding space, we apply our axes-based method and the classic gene-centric approach (detailed in Methods, section Assessing the ability of our axes-based methodology to capture biological information of the main manuscript and Supplementary Results, section Our axes-based method outperforms the classic gene-centric approach in capturing the cell's functional organization, respectively) and take the intersection between the GO BP terms captured by each method. We use these common GO BP terms to evaluate the agreement between the two approaches. The first column, "Embedding algorithm," lists the embedding algorithms used for generating the embedding spaces. The second column, "Rand," shows the adjusted Rand Index score between the clustering of these common GO BP terms averaged across dimensions and species. The third column, "SS Axes," displays Lin's semantic similarity between GO BP terms associated with the same axis averaged across dimensions and species. The fourth column, "SS Clusters," shows Lin's semantic similarity between GO BP terms enriched in the same gene cluster averaged across dimensions and species.

| Embedding algorithm | AUROC |
| --- | --- |
| ONMTF | 0.90 |
| NMTF | 0.90 |
| Deepwalk | 0.91 |

**Supplementary Table 14**: Our axes-based method and the FMM-based approaches agree with the biological information captured from different species PPI network embedding spaces. We generate the species PPI network embedding spaces by applying ONMTF, NMTF, and Deepwalk algorithms on the species PPI network of *Homo sapiens sapiens*, *Saccharomyces cerevisiae*, *Schizosaccharomyces pombe*, *Rattus norvegicus*, *Drosophila melanogaster*, and *Mus musculus* (detailed in Methods, sections Biological datasets and Embedding the PPI networks of the main manuscript). We generate these embedding spaces with increasing dimensionalities (from 50 to 1000 dimensions with a step of 50). For each species embedding space, we apply our axes-based method and our previous FMM-based approach [9] to uncover biological information from the spaces. Finally, we evaluate the agreement between the methods by investigating the functional interactions between the GO BP terms that they capture (detailed in Supplementary Results, section Our axes-based method is in agreement with the FMM-based methodology). The first column, "Embedding algorithm," lists the embedding algorithms used for generating the embedding spaces. The second column, "AUROC," displays the area under the receiver operating characteristic curve (AUROC) computed as detailed in Supplementary Results, Our axes-based method is in agreement with the FMM-based methodology.

[14] Howie D, Ten Bokum A, Necula AS, Cobbold SP, Waldmann H. The role of lipid metabolism in T lymphocyte differentiation and survival. Frontiers in

immunology. 2018;8:1949.

[28] Wang ZJ, Guan QW, Zhou HH, Mao XY, Chen FH. Mechanistic insight into lysyl oxidase in vascular remodeling and angiogenesis. Genes & Diseases. 2022;.
